# Structure of the human clamp loader bound to the sliding clamp: a further twist on AAA+ mechanism

**DOI:** 10.1101/2020.02.18.953257

**Authors:** Christl Gaubitz, Xingchen Liu, Joseph Magrino, Nicholas P. Stone, Jacob Landeck, Mark Hedglin, Brian A. Kelch

## Abstract

DNA replication requires the sliding clamp, a ring-shaped protein complex that encircles DNA, where it acts as an essential cofactor for DNA polymerases and other proteins. The sliding clamp needs to be actively opened and installed onto DNA by a clamp loader ATPase of the AAA+ family. The human clamp loader Replication Factor C (RFC) and sliding clamp PCNA are both essential and play critical roles in several diseases. Despite decades of study, no structure of human RFC has been resolved. Here, we report the structure of human RFC bound to PCNA by cryo-EM to an overall resolution of ~3.4 Å. The active sites of RFC are fully bound to ATP analogs, which is expected to induce opening of the sliding clamp. However, we observe the complex in a conformation prior to PCNA opening, with the clamp loader ATPase modules forming an over-twisted spiral that is incapable of binding DNA or hydrolyzing ATP. The autoinhibited conformation observed here has many similarities to a previous yeast RFC:PCNA crystal structure, suggesting that eukaryotic clamp loaders adopt a similar autoinhibited state early on in clamp loading. Our results point to a ‘Limited Change/Induced Fit’ mechanism in which the clamp first opens, followed by DNA binding inducing opening of the loader to release auto-inhibition. The proposed change from an over-twisted to an active conformation reveals a novel regulatory mechanism for AAA+ ATPases. Finally, our structural analysis of disease mutations leads to a mechanistic explanation for the role of RFC in human health.

## INTRODUCTION

DNA replication in all cellular life requires sliding clamps, ring-shaped protein complexes that wrap around DNA to topologically link numerous factors to DNA. Sliding clamps are necessary for DNA synthesis by DNA replicases, because they increase polymerase processivity and speed by orders of magnitude (Fay et al. 1981; Huang et al. 1981; Langston and O’Donnell 2008; Maki and Kornberg 1988; Mondol et al. 2019; Stodola and Burgers 2016). Sliding clamps additionally bind and facilitate function of scores of other proteins involved in diverse DNA transactions, such as DNA repair, recombination, and chromatin structure (Moldovan et al. 2007). The sliding clamp of eukaryotes, Proliferating Cell Nuclear Antigen (PCNA), is a critical factor for human health. PCNA’s central role in controlling many cancer pathways makes it a common cancer marker (Wang 2014). Recently, the genetic disease <u>P</u>CNA-<u>A</u>ssociated DNA <u>R</u>epair Disorder (PARD) was shown to be caused a hypomorphic mutation in PCNA that disrupts partner binding (Duffy et al. 2016; Baple et al. 2014).

PCNA’s ring shape necessitates active loading onto DNA by the RFC sliding clamp loader. Clamp loaders are five-subunit ATPase machines that can open the sliding clamp, and close it around DNA. Clamp loaders are found in all life, although their composition varies across different kingdoms (Kelch et al. 2012). The primary clamp loader in eukaryotes consists of five distinct proteins, RFC1 through RFC5. In humans, RFC plays a role in several diseases, such as cancer (Bell et al. 2011; von Deimling et al. 1999; Li et al. 2018), Warsaw Breakage Syndrome (Abe et al. 2018), CANVAS disease (Cortese et al. 2019), Hutchinson Gilford Progeria Syndrome (Tang et al. 2012), and in the replication of some viruses (Sun et al. 2014; Wang et al. 2012, 2013). However, it remains unknown whether loading by RFC contributes to the disease PARD.

Clamp loaders across all life appear to broadly share a similar mechanism. Clamp loaders are members of the AAA+ family of ATPases (<u>A</u>TPases <u>A</u>ssociated with various cellular <u>A</u>ctivities), a large protein family that uses the chemical energy of ATP to generate mechanical force (Erzberger and Berger 2006). Most AAA+ proteins form hexameric motors that use an undulating spiral staircase mechanism to processively translocate a substrate through the motor pore (Puchades et al. 2020; Gates et al. 2017; Puchades et al. 2017). Unlike most other AAA+ proteins, clamp loaders do not use ATP hydrolysis as a force generation step. Instead, the ATP-bound clamp loader forces the sliding clamp ring to open through binding energy alone (Xavier et al. 2000; Turner et al. 1999; Pietroni et al. 1997). Subsequent binding of primer-template DNA activates ATP hydrolysis, which results in clamp closure and ejection of the clamp loader (Ason et al. 2003; Berdis and Benkovic 1996; Gomes et al. 2001; Jarvis et al. 1989). Thus, the clamp loader is an ATP-dependent protein-remodeling switch.

The pentameric clamp loader structure is broadly conserved. The five subunits are named A through E going counter-clockwise around the assembly. Each of the five subunits consists of an N-terminal AAA+ ATPase module, followed by an α-helical ‘collar’ domain that serves to oligomerize the complex (Figure 1A). The Rossman fold and Lid domains that comprise the AAA+ module contain the catalytic residues for ATPase activity. Although most of the catalytic machinery is used *in cis*, the B, C, D, and E subunits all contain arginine finger residues that are provided *in trans* to complete the active site of a neighboring subunit.

**Figure 1.**
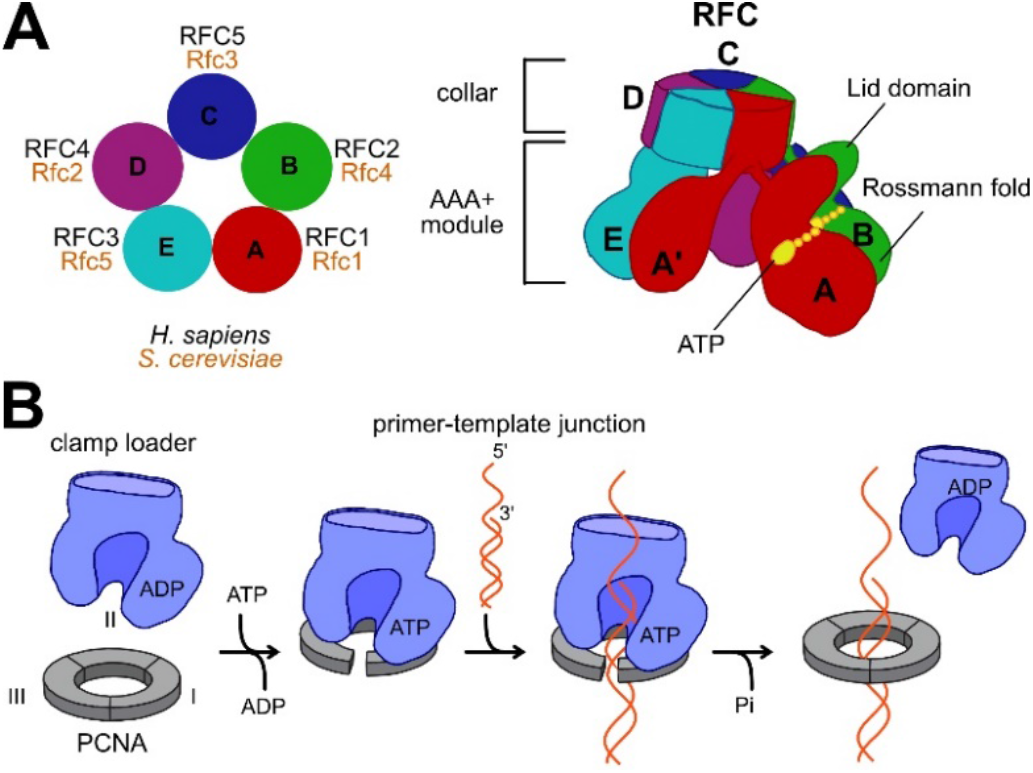
Human clamp loader (hRFC) composition and function. **(A)** hRFC consists of five different AAA+ ATPase subunits, named A to E. Each subunit consists of an ATPase module and a collar domain. The ATPase module has the active site sandwiched between an N-terminal Rossman fold ATPase domain and the Lid domain. **(B)** The clamp loading reaction begins with binding of ATP to the clamp loader, followed by clamp binding and opening. The clamp:clamp loader complex binds primer-template DNA, which triggers ATP hydrolysis, clamp closure, and clamp loader ejection.

We recently proposed a general mechanism for clamp loaders (Figure 1B) (Kelch 2016). The clamp loader, when bound to ATP, binds to and subsequently opens the sliding clamp (Gomes et al. 2001; Pietroni et al. 1997; Turner et al. 1999). Upon clamp opening, the complex binds to primer-template DNA inside the clamp loader’s central chamber (Chen et al. 2009; Goedken et al. 2004). DNA binding then activates ATP hydrolysis in the AAA+ active sites of the clamp loader, which results in clamp closure around DNA and ejection of the clamp loader (Ason et al. 2003; Berdis and Benkovic 1996; Gomes et al. 2001; Jarvis et al. 1989). The sliding clamp is now loaded at a primer-template junction for use by a DNA polymerase or other DNA metabolic enzymes.

Structural studies have revealed critical intermediates for the clamp loading mechanism. Early structures of the *E. coli* clamp loader in inactive states revealed the general organization of the complex (Davey et al. 2002; Kazmirski et al. 2004). A subsequent structure of a mutated form of the *S. cerevisiae* clamp loader RFC bound to PCNA showed RFC in a collapsed and over-twisted spiral conformation that is bound to a closed PCNA ring (Bowman et al. 2004). This conformation was initially hypothesized to represent an intermediate towards the end of the clamp loading reaction, with PCNA closed around DNA and still bound to RFC prior to ATP hydrolysis. It has also been hypothesized that this conformation is an artifact of the mutation of the arginine fingers that prevents proper assembly (Kelch 2016; Sakato et al. Hingorani 2012). The structure of an off-pathway intermediate of the *E. coli* clamp loader bound to a primer-template junction but no clamp confirmed the ‘notched-screwcap’ mode of DNA binding (Simonetta et al. 2009). Finally, the T4 phage clamp loader was crystallized with sliding clamp and DNA, revealing how ATP hydrolysis is linked to clamp closure (Kelch et al. 2011).

Despite many years of study, several central questions of clamp loader mechanism remain unanswered. Primarily, the mechanism of clamp opening is still unknown. It has been postulated that clamp opening occurs in a multi-step process, with initial clamp loader binding, followed by clamp opening (Chen et al. 2009; Goedken et al. 2004). However, there is no structure of this pre-opening intermediate state. Moreover, it remains unknown how disease mutations perturb clamp loader function, as there is currently no structure of the human RFC complex.

Here we describe a cryo-EM reconstruction of human RFC (hRFC) bound to PCNA. The structure reveals that PCNA is closed, despite all active sites of hRFC being bound to ATP analogs. The spiral of AAA+ modules is constricted, which prevents opening of the clamp and blocks the DNA-binding region in the central chamber of the clamp loader. We propose that this represents an autoinhibited form of the clamp loader that occurs prior to clamp opening. Our work provides a framework for understanding the clamp loader’s mechanism and function in human health.

## RESULTS

### Structure determination of the hRFC:PCNA complex

We sought to obtain a structure of human RFC bound to PCNA by single particle cryo-EM. We purified a hRFC construct with a truncation of the A subunit’s N-terminal region (RFC1ΔN555, missing residues 1-555) that expresses in *E. coli* (Supplemental Figure 1 A-B; (Kadyrov et al. 2009)). As expected (Uhlmann et al. 1997; Hedglin et al. 2013), our purified preparation of hRFC has highest ATPase activity in the presence of both the sliding clamp and primer-template DNA (Supplemental Figure 1C-D). For the rest of the paper, we refer to this complex as hRFC.

In order to visualize how the clamp loader interacts with the sliding clamp, we formed a complex of hRFC with PCNA and the slowly hydrolyzing ATP analog ATPγS. Brief cross-linking of the complex with bis(sulfosuccinimidyl)suberate (BS3) was necessary to obtain a quality reconstruction (Supplemental Figure 2A). BS3, which cross-links primary amines with a linker arm of ~11.4 Å, is used frequently to obtain high-resolution cryo-EM structures of labile complexes (Gerlach et al. 2018; Yoo et al. 2018). This treatment preserves the structure (Rozbeský et al. 2018) and mitigates issues with particle preferred orientation (Supplemental Figure 3B and Supplemental Figure 4A). Mass spectrometry reveals that most cross-links are intramolecular and ~55% of all crosslinks are in flexible loops that are not visible in the structure, with few cross-links connecting RFC to the clamp (Supplemental Figure 2B-C; Supplemental Table 1).

We determined the structure of hRFC:PCNA using single-particle cryo-EM. Representative images taken using a Titan Krios/K3 instrument are shown in Supplemental Figure 3A. We determined the cryo-EM structure to 3.4 Å using 3D classification and contrast transfer function (CTF) refinement (Supplemental Figure 3D). We improved the resolution of PCNA using multibody refinement with two masks covering hRFC and PCNA separately. The resolution of the PCNA-containing mask improved slightly to ~3.3 Å using the gold-standard criterion (FSC = 0.143; Supplementary Figure 3D and Supplemental Figure 4B). The local resolution ranges from ~2.9-4.1 Å with the highest resolution in the inner chamber of hRFC subunits A, B, and C, and the lowest resolution in peripheral loops (Supplemental Figure 4D). The resulting reconstruction was of sufficient quality to build an atomic model of hRFC (Supplemental Figure 4C). The high quality of the map allowed us to unambiguously assign each of the five chains to the respective gene product: RFC1 is the A subunit, RFC2 is B, RFC5 is C, RFC4 is D, and RFC3 is E (Figure 1A). Most parts of the protein could be easily modeled into the map, and our final model refines to an overall model-to-map correlation coefficient of 0.85 with good stereochemistry (Supplemental Table 2).

### The hRFC:PCNA complex is closed

Our reconstruction shows hRFC bound to a closed PCNA ring (Figure 2A-C). PCNA is in a planar, undistorted conformation, with only localized conformational changes where the ring is contacted by hRFC (Overall C_α_ RMSD is 0.98 Å; Supplemental Figure 5A). The closed PCNA ring was unexpected because clamp loaders typically open the sliding clamp when bound to ATP or ATP analogs (Hingorani and O’Donnell 1998).

**Figure 2.**
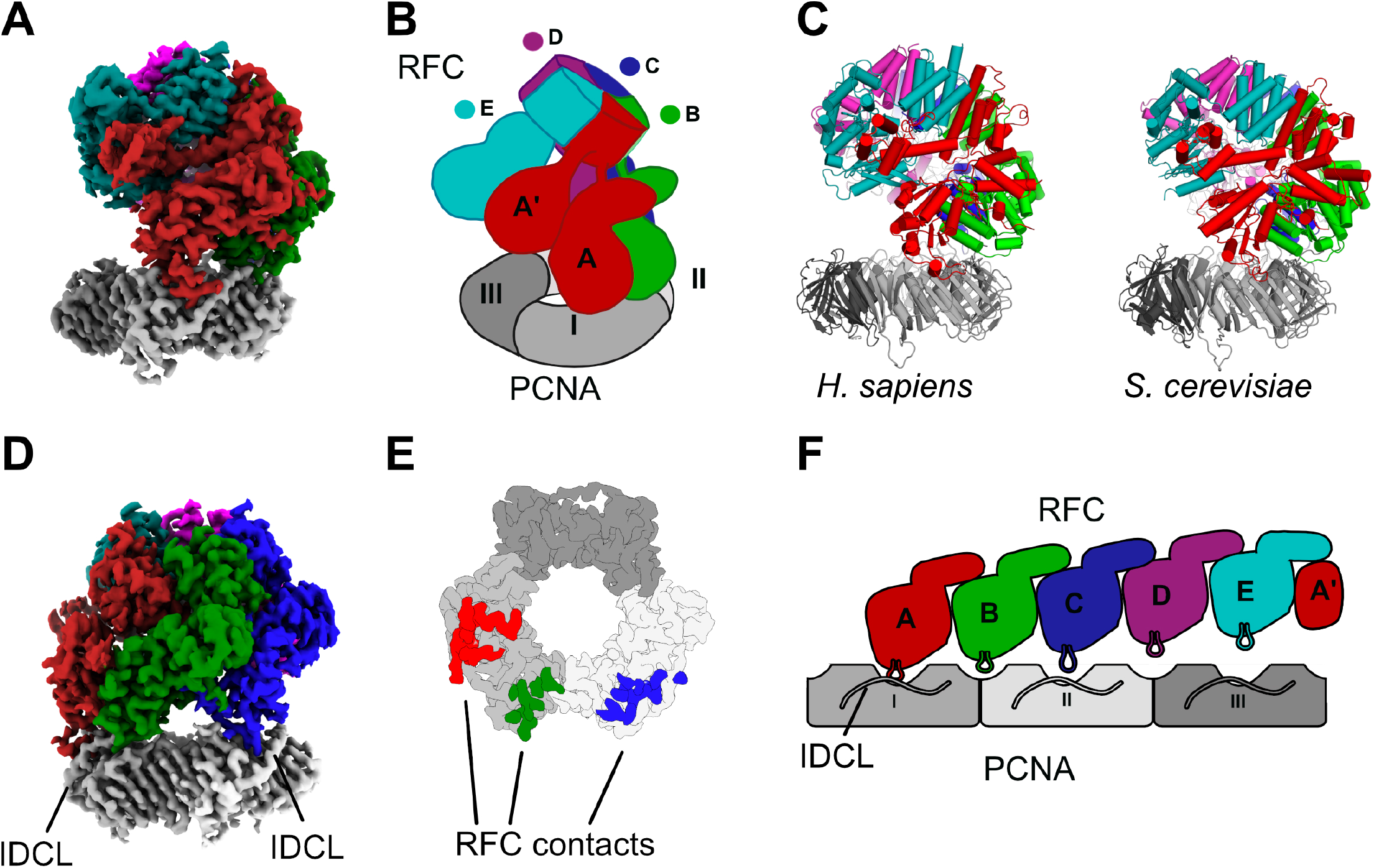
Architecture of hRFC bound to a closed PCNA ring. **(A)** The human hRFC:PCNA cryo-EM map is segmented and colored to show each subunit. **(B)** A cartoon presentation highlights that PCNA is closed. **(C)** Side view of the atomic model of hRFC:PCNA and the yeast RFC:PCNA crystal structure (PDB 1SXJ). **(D)** Back view of the hRFC:PCNA cryo-EM map showing the interaction sites of the B and C subunit with PCNA. **(E)** Top view on the interaction sites of the hRFC A, B and C subunits with PCNA. **(F)** Cartoon representation of the interaction sites of RFC and PCNA observed in this structure, with the PCNA and RFC AAA+ ATPase modules flattened onto the page. The A and C subunit contact the IDCL, whereas B is bound at the interface of PCNA subunits I and II. Subunits D and E do not contact PCNA.

Only the A, B and C subunits of hRFC contact the sliding clamp; the D and E subunits are lifted off the PCNA surface (Figure 2D-F). The A and C subunits bind PCNA’s <u>i</u>nter-<u>d</u>omain <u>c</u>onnecting <u>l</u>oop (IDCL), the primary binding site for PCNA interaction partners (Moldovan et al. 2007). The interaction between the A subunit of hRFC (hRFC-A) and subunit I of PCNA is the most extensive, burying ~2000 Å^2^ of surface area, much of this through hydrophobic interactions. The A subunit interacts with PCNA using a classic PIP-box motif that is similar in sequence and structure to the p21 protein, a high-affinity PCNA partner (Gulbis et al. 1996) (Supplemental Figure 5A and Supplemental Figure 6). The interaction between hRFC-C and the IDCL of PCNA subunit II is less extensive (~1200 Å^2^ of surface area buried) and does not induce the ‘high-affinity’ PCNA conformation (Supplemental Figure 5). Similar to hRFC-A, hRFC-C inserts an aromatic residue into the PIP-box binding cleft of PCNA to serve as the anchor (Supplemental Figure 5B). The RFC-B subunit binds the top of the interface between PCNA subunits I and II (Figure 2D-F). The hRFC-B:PCNA interaction is much less substantial than the other two sites, with only ~900 Å^2^ of buried surface area, primarily mediated by salt bridges. Therefore, the three contacts are not equal, with the hRFC-A interactions the strongest and the hRFC-B weakest.

### The AAA+ modules form an inactive, asymmetric spiral

All five subunits contain a bound nucleotide at the interfaces between hRFC subunits. The map is most consistent with ATPγS bound in the A, B, C, and D subunits, while the density in the E subunit is consistent with ADP, with no density at the γ-phosphate position (Figure 3A). The E subunit is inactive due to absence of several catalytic residues, so the presence of ADP cannot be due to phosphorothioate hydrolysis in the E site, but instead is likely from the ~10% contamination of ADP in typical preparations of ATPγS. The yeast and T4 phage clamp loaders also retain ADP in the inactive E subunit (Bowman et al. 2004; Kelch et al. 2011). Therefore, we hypothesize that ADP binding is a conserved function of the E subunit, and likely plays a structural role to stabilize this subunit. Related AAA+ machines such as the Origin Recognition Complex are thought to use nucleotide binding in a similar structural role (Bleichert 2019).

**Figure 3.**
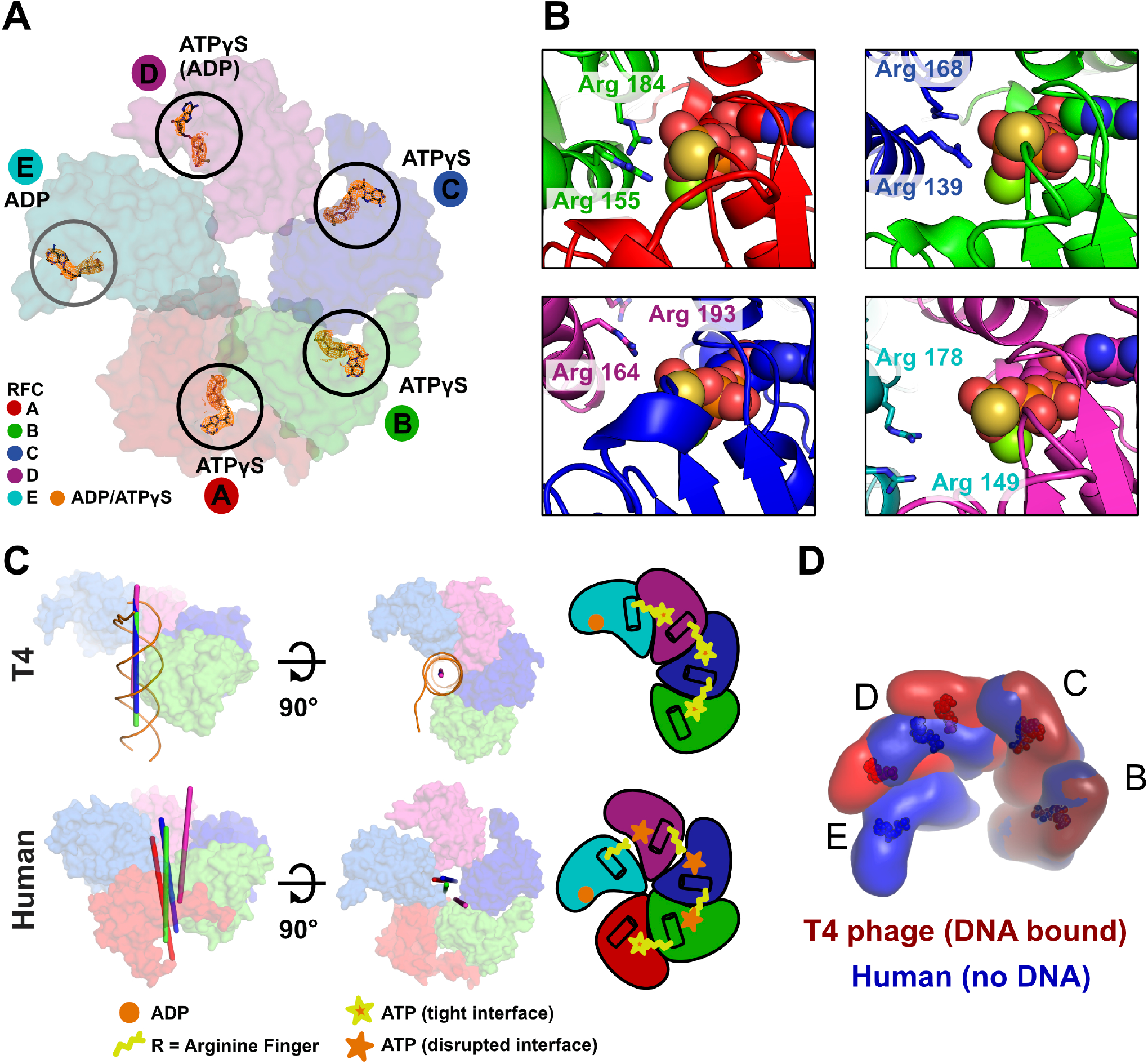
Architecture of the AAA-ATP domains and nucleotide binding. **(A)** Top view on the AAA+ domains of hRFC. All active sites (A, B, C, & D subunits) contain ATPγS. ADP is bound in the inactive E subunit. The nucleotide density at each site is shown in orange mesh. **(B)** Detailed view of all four active sites. The arginine fingers are distant in the B, C, and D ATPase sites, rendering them inactive. **(C)** Side and top views of the AAA+ spirals of the DNA-bound T4 phage clamp loader (PDB 3U60, top panel), and hRFC. The rotation axes that relate the A to B, the B to C, the C to D, and the D to E subunits are shown in red, green, blue, and magenta, respectively. The rotation axes of T4:DNA complex are coincident with each other and the central axis of DNA; in contrast, the axes of hRFC are severely skewed. **(D)** The AAA+ spiral of hRFC is over-twisted relative to the active, DNA-bound form.

Most active sites are not in an active conformation. The interfaces for the B, C, and D active sites are disrupted, such that the *trans*-acting arginine fingers are not in a productive conformation (Figure 3B). The B-C interface is tighter than the C-D and D-E interfaces but is still too expanded for contact by the arginine fingers to γ-phosphorothioate (~ 4 Å and 11 Å for the SRC motif arginine). The D subunit is swung out, which even further disrupts contacts between arginine fingers to the active sites at the C and D subunits (~7 Å and ~17 Å for Arg164 and Arg193 of hRFC-D; ~9 Å and ~8 Å for Arg149 and Arg178 of hRFC-E) (Figure 3B). The interface between the A and B subunits is tightest and similar to that found in active conformations found in other clamp loaders (Kelch et al. 2011; Simonetta et al. 2009) (Figure 3A-B). However, this active site is unnecessary for clamp loading in all loaders tested (Schmidt et al. 2001; Sakato et al. 2012; Seybert and Wigley 2004), so the functional relevance of this conformation is unclear. Therefore, all ATPase sites required for clamp loading activity are in an inactive conformation.

The hRFC AAA+ spiral is asymmetric and over-twisted, which sterically hinders DNA binding (Figure 4A). The axes of rotation relating adjacent AAA+ domains are not coincident with each other, indicating that the spiral in hRFC lacks helical symmetry (Figure 3C). The hRFC spiral observed here has a reduced helical radius, although the pitch is similar to that of the DNA bound spirals. In contrast, the structures of T4 and *E. coli* clamp loaders bound to DNA show a symmetric spiral of AAA+ domains around DNA (Figure 3C and Supplemental Figure 7) (Simonetta et al. 2009; Kelch et al. 2011). The D and E subunits of hRFC are particularly over-twisted such that they fill the DNA binding region (Figure 3D and Figure 4A). Moreover, the intramolecular interaction between the A’ domain and the A subunit AAA+ module would prevent DNA access to the central chamber of hRFC. The interaction between the two domains are rather weak with little buried surface area (~250 Å^2^) and poorly resolved, and is therefore unlikely to be the main driving force for over-twisting the AAA+ spiral. The over-twisted state of the AAA+ spiral that we observe for hRFC has important ramifications for how hRFC opens PCNA and binds DNA (see Discussion).

**Figure 4.**
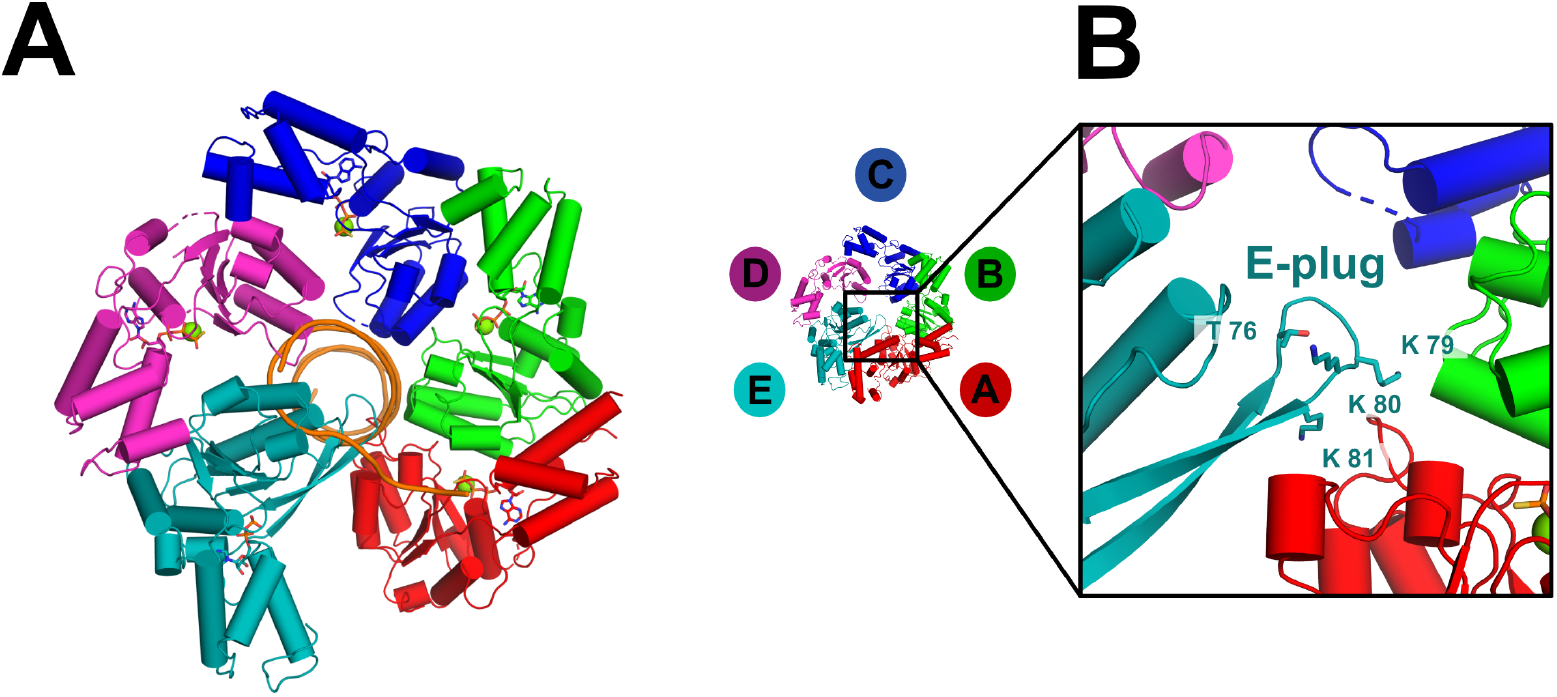
The E-plug blocks the DNA binding chamber. **(A)** The hRFC AAA+ spiral (top view) is incompatible with DNA binding, primarily through clashes with the D and E subunits. DNA is superposed from the structure of T4 phage clamp loader bound to DNA (PDB 3U60). **(B)** The ‘E-plug’ is a β-hairpin that extends into the DNA binding chamber. The tip of the E-plug contains conserved basic residues.

The conformation of the hRFC:PCNA complex we observe here is very similar to the crystal structure of the yeast RFC:PCNA complex (Figure 2C and Supplemental Figure 7A-B) (Bowman et al. 2004). Both structures contain a closed PCNA ring contacted by only the A, B, and C RFC subunits, with similar buried surface between the two proteins (4600 vs. 5200 Å^2^ buried surface area for human and yeast RFC:PCNA complexes respectively). The AAA+ spirals of hRFC and yRFC adopt similar conformations, with only a small difference in position of the D subunit (Supplemental Figure 7B).

### The E-plug region fills the DNA binding chamber

We observe a β-hairpin of the hRFC E subunit that blocks the DNA binding region (Figure 4A-B). This β-hairpin extends the β-sheet in the Rossman fold domain of the AAA+ module, forming a mixed parallel/antiparallel β-sheet. To our knowledge, this type of topology is not found in other AAA+ proteins. We call this β-hairpin the ‘E-plug’ because it inserts into and completely blocks the DNA binding region by interacting with the Rossman fold of the A subunit and possibly the B subunit (Figure 4A and Supplemental Figure 8B). The E-plug was not modeled in the yeast RFC crystal structure, presumably because of weak density. However, sequence alignments reveal that the E-plug feature is conserved in RFC-E subunits throughout eukaryotes (Supplemental Figure 8A). There are conserved basic residues at the tip of the β-hairpin (lysine 79-81, Figure 4B, Supplemental Figure 8A). We hypothesized that the E-plug changes its position to interact with the DNA backbone upon DNA binding. Moreover, there are two known phosphorylation sites at the tip of the E-plug: Thr75 and Thr76. Phosphorylation of Thr76 is thought to be important because this phosphosite has a high ‘Functional Score’, a quantitative prediction of phosphosite importance based on aggregation of nearly 60 different features (Ochoa et al. 2019). Therefore, the E-plug appears to be important for clamp loader function and may act as a site of clamp loader regulation.

We hypothesized that the E-plug controls RFC function. Because the E-plug tip is conserved to retain positive charge, we hypothesized that these residues are important for binding DNA. Furthermore, we hypothesized that phosphorylation of Thr76 would inhibit DNA binding. To test these hypotheses, we constructed two hRFC variants: one in which the three lysines at the E-plug tip were simultaneously mutated to alanine (K79A, K80A, K81A or the 3K→3A mutant), and the phosphomimetic mutant Thr76Asp (T76D). To determine if the mutations perturb clamp loader function, we measured the PCNA- and DNA-dependency on ATPase activity for each of these variant hRFC complexes. We find that hRFC-3K→3A exhibits ~2-fold lower maximal ATPase activity than wild-type hRFC (Supplemental Figure 8C). In contrast, the ATPase activity of hRFC-T76D is nearly the same as WT-hRFC. We then measured ATPase activity as a function of DNA concentration, which we use to estimate DNA binding affinity (Supplemental Figure 8C-D). The DNA dependence of ATPase activity reveals that both variants bind DNA with equivalent affinity as WT-hRFC. These results suggest that the E-plug plays a role in ATPase activation, but is not critical for DNA binding affinity.

### Investigating disease mutations in hRFC and PCNA

To investigate how RFC mutations commonly found in cancer could affect loading of PCNA, we mapped somatic cell cancer mutations onto the hRFC:PCNA structure using the Catalogue of Somatic Mutations in Cancer (COSMIC) database. To enrich for putative driver mutations, we focused on the most common mutation sites (analyzed sites have missense and nonsense mutations from three or more patient isolates; Supplementary Table 3). Furthermore, we analyzed these mutations with Rhapsody, a computational tool that estimates the pathogenicity of missense mutations (Ponzoni et al. 2020).

Cancer mutations are significantly more prevalent in the collar domain than in the AAA+ spiral or the sliding clamp (P_value_ = 0.006) (Figure 5A and Supplemental Figure 9A). Of the most common 24 mutations in RFC or PCNA, 42% are in the collar region, even though collar domains account for only ~18% of the total length of the combined clamp loader subunits. In contrast, the AAA+ spiral region (AAA+ modules plus the A’ domain) harbors 42% of the mutations, but it accounts for 58% of the total clamp loader length. We also note a slight preference for mutations to occur in the D subunit over other RFC subunits, although this trend is not statistically significant (P=0.1). The D subunit (RFC4) has a cancer mutation hit rate twice the average of all other RFC subunits (0.017 vs. 0.008 mutations per residue, for the D subunit vs. all other subunits. Therefore, we find that the collar region and, to a lesser extent, the D subunit tend to be hotspots for cancer mutations. Additionally, we also note that PCNA and the N-terminal and C-terminal regions of the A-subunit have particularly low cancer mutation hit rates. There are no mutation sites in PCNA with a hit-count of three or higher, and only a handful of residues with a hit-count of two. Furthermore, Rhapsody predicts that each of the five RFC subunits contains at least one deleterious missense mutation (Supplemental Figure 9B and 9C).

**Figure 5.**
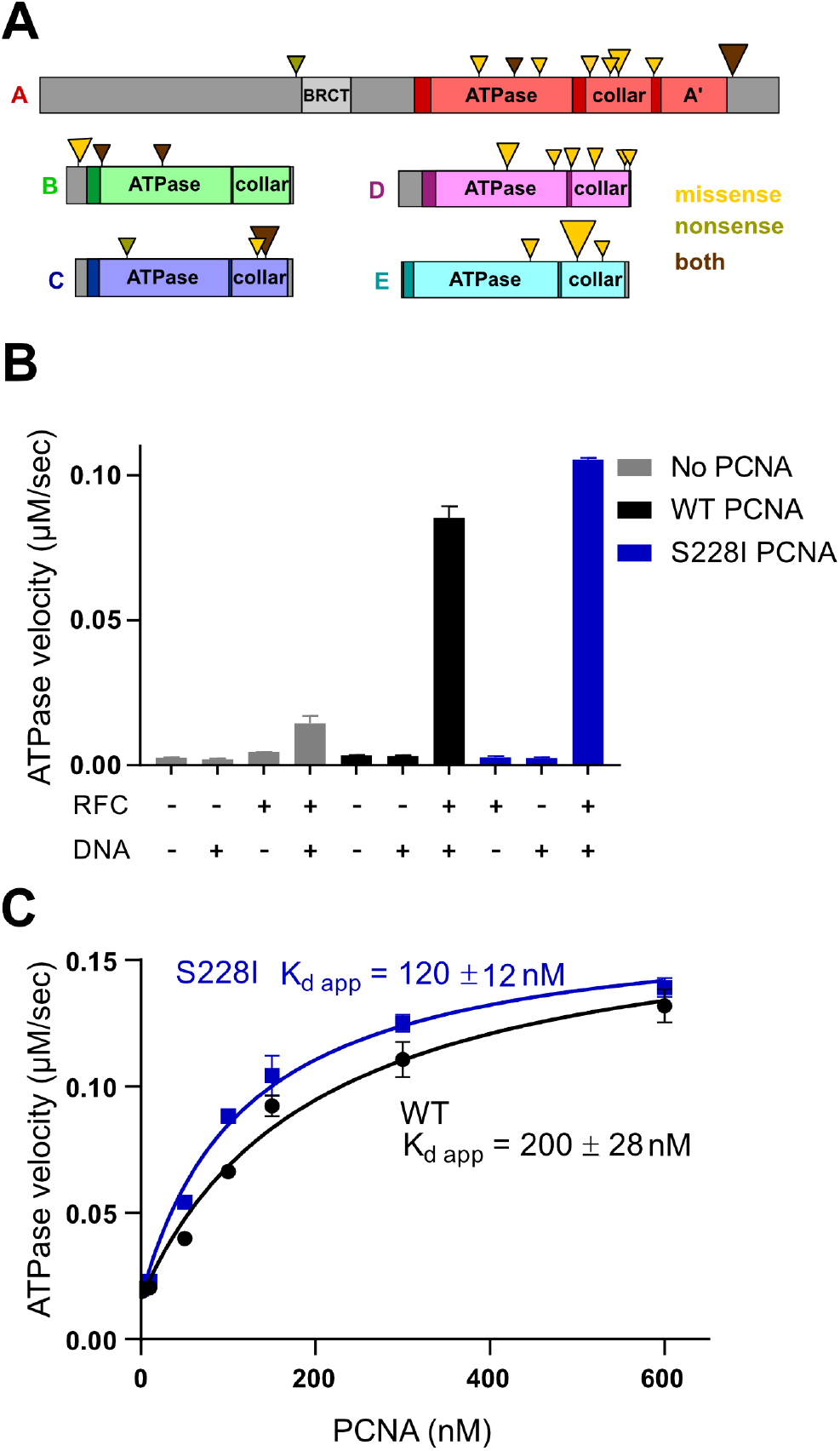
hRFC in disease. **(A)** Cancer mutations mapped onto primary sequence of hRFC. Triangle size is scaled by the number of hits at that site and colored according to the mutation type. Gray regions are not visible in the cryo-EM structure. **(B)** hRFC ATPase activity dependence on WT-PCNA and S228I-PCNA. S228I-PCNA has a minor effect on the ATPase hydrolysis rate of hRFC. **(C)** Rate of hRFC ATP hydrolysis as a function of WT-PCNA and S228I-PCNA concentration. S228I-PCNA has a mild enhancement in hRFC affinity.

Our structure also sheds light on a rare genetic disease PARD, which is caused by a hypomorphic mutation in PCNA that converts serine 228 to isoleucine (Baple et al. 2014). We had previously shown that the S228I mutation deforms the IDCL of PCNA, thereby weakening the affinity of some partners (Duffy et al. 2016). However, a major question remained: does the S228I mutation perturb PCNA loading? A simplistic structural analysis would suggest that the interaction between hRFC and S228I-PCNA is disfavored (Supplemental Figure 5B). To test this hypothesis, we measured hRFC ATPase activity as a function of PCNA concentration, which gives an estimate of the affinity for PCNA (Figure 5 B-C). We find that the S228I-PCNA mutation has no effect on maximal activity, (V_max,WT_ = 0.16 ± 0.01 *μ*M/sec, V_max,S228I_ = 0.15 ± 0.01 *μ*M/sec) and only a mild stimulatory effect on affinity (apparent K_d_ of 200 ± 28 nM and 120 ± 12 nM for WT and S228I, respectively). In contrast to expectations, these results suggest that the disease mutation causes no substantial defect on PCNA loading.

## DISCUSSION

### Implications for clamp loading

We were expecting to obtain a structure of hRFC bound to an open sliding clamp, but we did not observe a significant number of particles with PCNA in the open form. Previous FRET experiments indicate that PCNA is rapidly opened after initial binding to RFC (Zhou and Hingorani 2012; Trakselis et al. 2001; Perumal et al. 2019). Furthermore, our cross-linking experiments indicate strong cross-linking between Lys144 of the E subunit with Lys254 of PCNA, an interaction that can only occur once PCNA is opened (Kelch et al. 2011). Therefore, the open PCNA complex is likely populated in solution, but these open species are currently not suitable for high resolution structure determination. We hypothesize that the open PCNA complex interacts with the air-water interface, likely through exposed hydrophobic residues in the open clamp, which may result in aggregation or denaturation (D’Imprima et al. 2019; Noble et al. 2018).

The conformation of the hRFC:PCNA complex captured here is very similar to the crystal structure of a mutated form of the yeast RFC:PCNA complex (Bowman et al. 2004). This similarity is surprising, as it had been presumed that mutation of the arginine finger residues caused this ‘over-twisted’ state (Kelch 2016; Sakato et al. 2012). The structural similarity observed here using a functional version of the hRFC complex suggests that this state is actually well-populated in solution and is conserved across ~1 billion years of evolution since the *S. cerevisiae* and *H. sapiens* lineages diverged (Douzery et al. 2004). If so, what intermediate in clamp loading does it represent?

We propose that the conformation observed here represents the first encounter complex between RFC and PCNA. All four of the ATPase sites are occupied by ATP analog, indicating that no ATP hydrolysis is necessary for this state to accumulate. Because ATP hydrolysis occurs at the end of clamp loading (Kelch et al. 2012), our structure must represent a state at the beginning of the clamp loading cycle, before clamp opening. The Hingorani group has proposed that yeast RFC first binds to PCNA without ring opening (Liu et al. 2017); we propose that the conformation captured here by cryo-EM is that first encounter complex (Figure 6).

**Figure 6.**
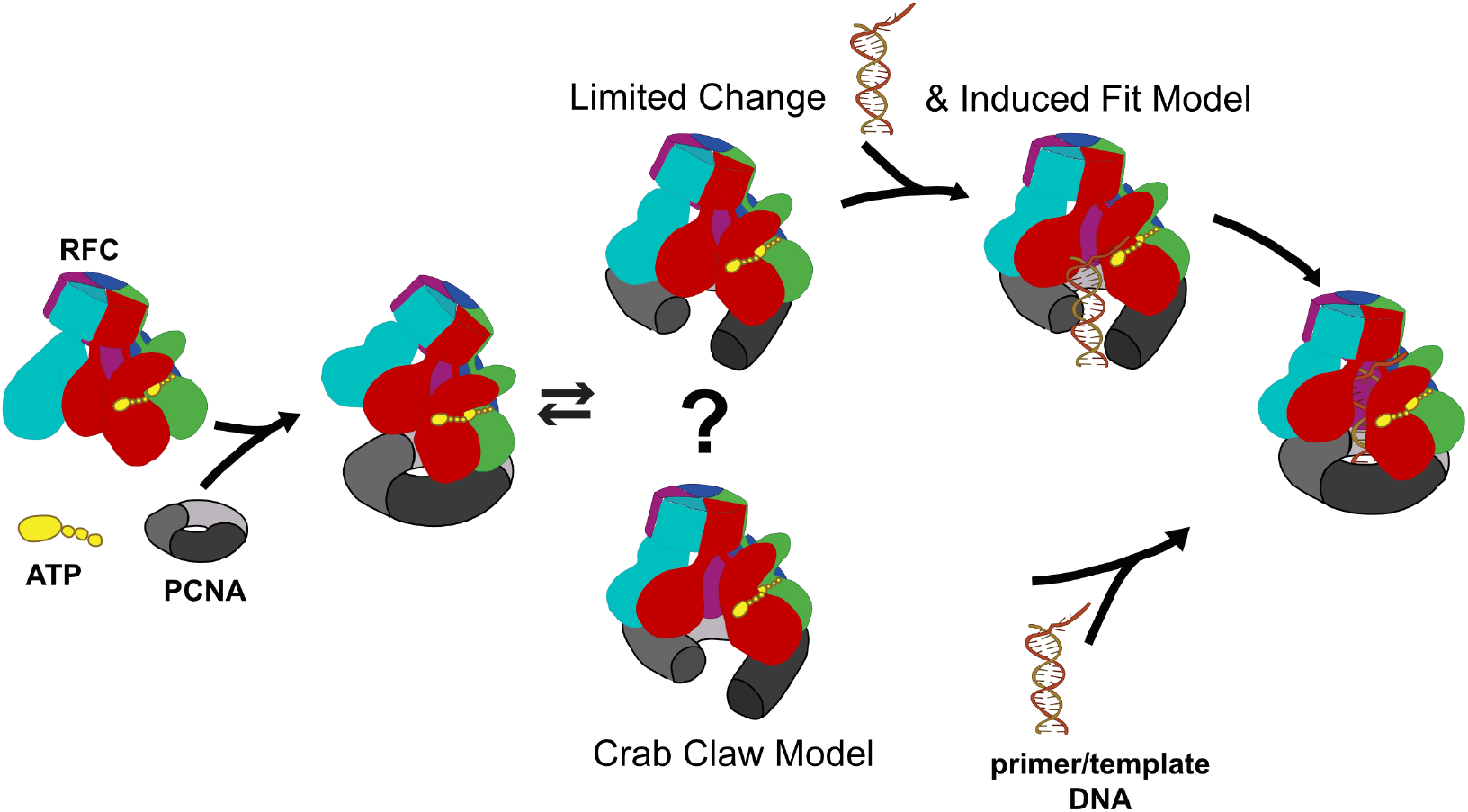
Model for PCNA opening. In the limited change model, PCNA is opened without RFC undergoing a large conformational change. Contact with primer-template-DNA induces a widening of RFC’s spiral to allow for binding into RFC’s central chamber. Alternatively, in the crab-claw model, RFC undergoes a large conformational change concurrent with clamp opening to open up the DNA binding chamber.

Our structure indicates that hRFC can adopt an autoinhibited conformation. The AAA+ spiral is distorted and over-twisted, which disrupts the interfacial contacts necessary for ATPase activity and blocks the inner DNA-binding chamber. Access to the DNA binding region is further blocked by the E-plug, which reaches across the central chamber of the clamp loader to make extensive contacts with the Rossman fold of the A subunit (Supplemental Figure 8B). Typically, the single-stranded region of template DNA extrudes from the gate between the A’ domain and the AAA+ domain (Kelch et al. 2011) (Simonetta et al. 2009). This interface is closed in our structure. Therefore, this state can neither bind DNA nor hydrolyze ATP. This autoinhibited conformation may limit wasteful hydrolysis of ATP by maintaining this inactive conformation. Only binding of both clamp and DNA places the clamp loader into an active ATPase conformation. This auto-inhibited state may also allow for clamp loader regulation through post-translational modification or binding of accessory factors.

We identified the E-plug as a novel structural element of the RFC-E subunit that blocks the DNA binding chamber. Based on sequence analysis, the E-plug is conserved in all known eukaryotic RFC-E subunits, yet we know of no analogous feature in other AAA+ proteins. The E-plug blocks DNA binding in the conformation observed here, but could be playing different roles when DNA is bound. The residues at the tip of the E-plug are conserved as positively charged, perhaps hinting at interactions with DNA (Supplemental Figure 8A, 8B and Figure 4B). Our data suggest that these residues are not important for DNA binding affinity, but could play a role in sensing the presence of DNA for ATPase activation (Supplemental Figure 8C and 8D).

The over-twist in the AAA+ spiral occurs primarily at the D subunit. The D subunit deviates from an active symmetric conformation more than any other subunit (Figure 3). This is recapitulated in the crystal structure of the yeast RFC arginine finger variant (Bowman et al. 2004), indicating that the D subunit’s unusual positioning is conserved. The placement of the D subunit’s AAA+ module disrupts two different interfaces: C-D and D-E. Therefore, we propose that the D subunit plays a key role in mediating autoinhibition. In support of this hypothesis, kinetic studies indicate that the C and D subunit’s ATPase activity dictates clamp loader function (Sakato et al. 2012). The entire clamp loader can be kept in an auto-inhibited state by inactivating the ATPase sites that control clamp loader activity. Taken together, the D subunit’s role is particularly ‘pivotal’ for RFC function: the D subunit is an essential component of the RFC machine, and it acts as a pivot point for AAA+ motion.

Based on our structure, we hypothesize that the clamp loader undergoes a conformational change that opens the gate between A’ and the AAA domain in A subunit to allow primer-template DNA binding within the RFC inner chamber. Previous Fluorescence Resonance Energy Transfer experiments of the *E. coli* clamp loader indicated that addition of ATP and/or the sliding clamp does not trigger opening of the gate region (Goedken et al., 2004). This study led to the Limited Change Model, which suggests that the ATP-bound clamp loader is already in a conformation competent to open the sliding clamp. Based on our data, we now modify this model to include the presence of this auto-inhibited state. In this Limited Change/Induced Fit model, clamp binding and opening occurs with minimal change in clamp loader conformation. Subsequent binding of primer-template-DNA within the open clamp induces a conformational change in the loader that opens the central chamber so that DNA can be productively bound by both loader and clamp (Figure 6). In support of this hypothesis, Molecular Dynamics simulations of PCNA opening with yeast RFC suggested that clamp opening is possible with limited conformational change in RFC (Tainer et al., 2010). Another possible mechanism, the Crab-Claw Model, posits that PCNA opening triggers a concomitant conformational change within RFC that opens the central chamber for DNA binding. However, the Crab claw mechanism does not agree with the previous fluorescence studies (Goedken et al. 2004). Future studies will differentiate between the two mechanisms for clamp opening.

How does the autoinhibited state we see with RFC compare with those of other AAA+ machines? The revolution in cryo-EM has led to the recent proposal that many AAA+ machines are processive motors that function using an undulating spiral staircase mechanism (Puchades et al. 2017; Puchades et al. 2020; Gates et al. 2017). However, our work here and elsewhere indicates that clamp loaders appear to use a different mechanism (Kelch et al. 2011; Kelch 2016). The inactive state reported here is very distinct from the active conformations observed when bound to DNA (Simonetta et al. 2009; Kelch et al. 2011). Therefore, the helical radius of the AAA+ spiral must expand to accept DNA, indicating that the primary mechanism of clamp loader activation is controlled by the shape of the AAA+ spiral. Other AAA+ switches such as the Origin Recognition Complex show auto-inhibition mechanisms wherein the spiral is disrupted, although the disruption is not due to over-twisting (Bleichert 2019). In contrast, in the ‘spiral staircase’ view of processive AAA+ motors, the helical radius is fairly uniform during function (Puchades et al. 2020). The over-twist of the AAA+ spiral may be due to the fact that RFC is pentameric and not hexameric. The lack of the sixth subunit may afford the space for the helical radius to significantly change, particularly in the absence of DNA. This regulatory mechanism may not be readily available for hexameric AAA+ motors because the sixth subunit creates a steric block. Therefore, the pentameric clamp loaders function in a very distinct manner from hexameric AAA+ machines, which allows for distinct modes of regulation.

### Implications for cancer and the rare diseases PARD and HGPS

We identified cancer mutation hotspot regions, with a strong preference for mutations occurring in the collar region (Figure 5A and Supplemental Figure 9A). Why is the collar a hot spot for mutation? The collar is absolutely critical for clamp loader assembly, so it stands to reason that many of the cancer mutations disrupt proper oligomerization of hRFC. We propose that improperly assembled clamp loaders are particularly deleterious to the cell. This is reflected by Rhapsody’s pathogenicity assignment, where the most common cancer mutation in hRFC at the A-E collar interface is predicted to be deleterious (Supplemental Figure 9B and 9C). An incomplete hRFC could possibly have dominant negative effects by inducing premature unloading, as has been seen *in vitro* for the bacterial clamp loader (Leu et al. 2000). Additionally, disruption of the oligomerization interfaces may alter the relative populations of hRFC and the three alternative clamp loaders, which are important guardians of genome integrity (Majka and Burgers 2004). We also identified a weak preference for cancer mutations to accumulate in the D subunit. The D subunit is also most often found to be amplified in various cancers (Li et al. 2018). As noted above, the D subunit is particularly important for over-twisting the AAA+ spiral into the auto-inhibited conformation. Perhaps this specialized role of the D subunit makes it more likely to accumulate mutations that can drive cancer.

Our work has also revealed insights into PARD, a rare disease caused by a hypomorphic mutation of PCNA that results in misregulation of the sliding clamp (Duffy et al. 2016; Wilson et al. 2017; Baple et al. 2014). Because clamp loading is the primary means of PCNA regulation (Moldovan et al. 2007; Kelch 2016), understanding the mechanistic impacts of the PARD S288I mutation on clamp loading is of prime importance. We expected that loading of S228I-PCNA would be less efficient because our structure predicts that the interaction between hRFC and S228I-PCNA would be disfavored (Supplemental Figure 5B). However, we observe that the S228I mutation does not reduce hRFC ATPase activity or binding affinity (Figure 5B&C), indicating that loading of S228I-PCNA is unlikely to be a driver of PARD disease.

Our results provide insight into the mechanism by which S228I-PCNA interacts with its partners. S228I-PCNA causes a dramatic loss in affinity for partners that have typically bind PCNA with moderate to low affinity, such as FEN1 and RNaseH2 (Duffy et al. 2016; Wilson et al. 2017; Baple et al. 2014). However, our work here shows that tight-binding partners, such as hRFC and p21, maintain their tight binding affinity for S228I-PCNA. Therefore, we propose that absolute binding affinity determines whether the partner loses affinity for the disease mutant. In other words, we predict a correlation between the ΔG_binding_ and ΔΔG_WT-S228I_. Furthermore, we find that RFC-A binds PCNA using a PIP-box motif that closely resembles that of p21 (Supplemental Figure 6). Both p21 and hRFC-A have a tyrosine residue at the second conserved aromatic position of the PIP-box (Tyr151 and Tyr703 in p21 and hRFC-A, respectively), unlike the most partners that have phenylalanine at this position. We hypothesize that the hydroxyl group of Tyr703-RFC1 reorients the IDCL of S228I-PCNA into the tight binding configuration, as Tyr151 does for p21 (Duffy et al.2016). In support of our hypothesis, the presence of tyrosine at the second aromatic position of the PIP-box can increase affinity (Kroker and Bruning 2015).

Our structure also provides insight into Hutchinson-Gilford Progeria Syndrome (HGPS) because the hRFC construct that we used here is similar to the HGPS variant (Tang et al. 2012). Both the HGPS disease variant and the construct used in this study remove the N-terminal region of the A subunit RFC1, including the BRCT domain. The RFC A subunit (RFC1) is proteolytically truncated to a ~75 kDa C-terminal fragment in HGPS, removing approximately the first 500 residues, near the N-terminal end of a linker connecting the BRCT domain to the AAA+ module (Tang et al. 2012). The construct we used for structure determination is also truncated within this predicted linker region, at residue 555. The HGPS variant is thought to be a less effective clamp loader, because chromatin-bound PCNA levels drop as the levels of truncated clamp loader increase (Tang et al. 2012). Perhaps the N-terminal region assists in the clamp loading process. If so, this could explain why we failed to observe hRFC bound to an open PCNA; if this region is important for holding PCNA open, then the construct we used for structure determination may diminish the population of open hRFC:PCNA complexes. Although the RFC1 ΔN555 construct can load PCNA onto DNA (Uhlmann et al. 1997; Hedglin et al. 2013; Podust et al. 1998), it is unknown if the truncation affects clamp loading efficiency *in vivo*. The RFC1 N-terminal region has been hypothesized to assist in localizing RFC to DNA non-specifically (Podust et al. 1998), again suggesting that there could be a DNA binding defect in the HGPS variant. Future work will examine the potential linkage of the N-terminal region with clamp loader function and human disease.

## MATERIALS AND METHODS

### Protein expression and purification

p36-p37-p38-p40-pET-Duet-1 was modified using site-directed mutagenesis to clone the E-plug mutants (Rfc3-3KtoA and Rfc3-T76D) (Liu and Naismith 2008). hRFC was overexpressed and purified following the protocol of (Kadyrov et al. 2009; Perumal et al. 2019) with minor modifications. p36-p37-p38-p40-pET-Duet-1 and pCDF-1b-RFC140 plasmids were co-transformed into commercial BL21(DE3) *E. coli* cells (Millipore). After preculture, transformants were grown in 6 liters of prewarmed terrific broth medium supplemented with 50 μg/mL streptomycin and 100 μg/mL ampicillin at 37 °C and induced with IPTG at an optical density of 0.8. Protein expression was continued at 18 °C for 15 hours.

Cells were harvested by centrifugation at 7,277 g for 20 min, resuspended in 200 mL of 20 mM Hepes-KOH, pH 7.4 with 200 mM NaCl and pelleted by centrifugation at 4,000 g for 20 min. The pellet was resuspended in 15 mL lysis buffer per one cell optical density. The lysis buffer contained 20 mM HEPES KOH, pH 7.4, 180 mM NaCl, 2 mM EDTA, 5% glycerol (w/v), 0.01% NP-40 (v/v), 2 mM DTT, and Roche protease inhibitor mix. Cells were lysed using a cell disruptor, pelleted and the supernatant was filtered.

After filtration, the supernatant was applied to a 25 ml HiTrap SP HP column (GE Healthcare). The column was washed with three column volumes of SP column buffer (25 mM HEPES KOH, pH 7.4, 1180 mM NaCl, 0.1 mM EDTA, 5% glycerol (w/v), 0.01% NP-40 (v/v), 2 mM DTT) and developed with a 7 column volumes linear NaCl gradient (200–1,000 mM). Fractions containing all RFC subunits were diluted with 5% glycerol (w/v), 0.01% NP-40 (v/v), 50 mM KPO_4_ buffer, pH 7.5 to a salt concentration of 100 mM NaCl and loaded onto a 5 mL Bio-Scale TM Mini CHT Type II column (Bio-Rad) equilibrated with CHT column buffer (5% glycerol (w/v), 0.01% NP-40 (v/v), 50 mM KPO4 buffer, pH 7.5, 100 mM NaCl). After a 2.5 column volume (CV) wash, the protein was eluted with a stepwise gradient of 140 mM (3 CV), 185 mM (2.5 CV) 230 mM (2.5 CV), 275 mM (2.5 CV), 350 mM (2.5 CV), and 500 mM (2.5 CV) KPO_4_ buffer, pH 7.5. Peak fractions of hRFC were pooled, buffer exchanged and concentrated with an Amicon device to 15-20 mg/ml into a buffer containing 25 mM HEPES-KOH, pH 7.4, 15% glycerol (w/v), 0.01% NP-40, 300mM NaCl, 2 mM DTT for storage. The protein was further purified by gel filtration with a Superose 6 10/300 GL column (GE Healthcare). hPCNA was expressed and purified as described in (Duffy et al. 2016).

### ATPase enzymatic Assays

hRFC was incubated at room temperature with a master mix (3U/mL Pyruvate kinase, 3 U/mL Lactate dehydrogenase, 1 mM ATP 670 μM Phosphoenol pyruvate, 170 μM NADH, 50 mM Tris (pH 7.5), 500 μM TCEP, 5 mM MgCl_2_, and 200 mM Potassium glutamate) with 1 *μ*M PCNA and varying concentrations of annealed oligonucleotides. The annealed DNA has a 10-base 5’ overhanging end. The template strand sequence was 5’-TTTTTTTTTTTATGTACTCGTAGTGTCTGC-3' and the primer strand sequence 5’-GCAGACACTACGAGTACATA–3’ with a recessed 3’-end. ATPase activity of hRFC was measured in a 96-well format with a Perkin–Elmer Victor3 1420 multichannel counter using an excitation filter centered at 355 nm, with a bandpass of 400 nm to detect NADH oxidized to NAD+.

Initial rates were obtained from a linear fit of the initial slopes. Rates were plotted as a function of primer-template DNA concentration and the data was fit using a hyperbolic equation:

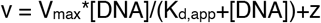

where V_max_ is the maximum enzyme velocity K_d,app_ is the substrate concentration needed to get a half-maximum enzyme velocity, and z is the velocity with no DNA (GraphPad Prism).

### Crosslinking and Mass Spectrometry

hRFC was crosslinked with the amine-reactive reagent Bis(sulfosuccinimidyl)suberate (BS3, Thermo Scientific Pierce). For crosslinking, hRFC and hPCNA were mixed in a in a ratio of 1/1.3 respectively and buffer exchanged using an Amicon Ultra-0.5 mL centrifugal concentrator (Millipore) into buffer containing 1 mM TCEP, 200 mM NaCl, 50 mM, Hepes-NaOH, pH 7.5, and 4 mM MgCl_2_. The protein was diluted to 3 μM and after the addition of 1 mM ATPγS and a wait time of 3 min, 1 mM of BS3 was added for crosslinking. The sample was incubated for 15 min at room temperature. The crosslinking reaction was neutralized with Tris-HCl. For mass spectrometry, the sample was snap frozen in liquid nitrogen and shipped on dry ice.

The purified complex was reduced, alkylated, and loaded onto an SDS-PAGE gel to enrich for the crosslinked complex by size. The gel >150kDa was excised, destained, and subjected to proteolytic digestion with trypsin. The resulting peptides were extracted and desalted as previously described (Peled et al. 2018) and an aliquot of the peptides was analyzed with LC-MS coupled to a ThermoFisher Scientific Q Exactive Mass Spectrometer operated in data dependent mode selecting only precursors of 3. The data was searched against the UniProt human database, using Byonic and XlinkX within Proteome Discoverer 2.3.

### Electron Microscopy

#### Negative-Staining EM

Purified hRFC was diluted to a concentration of 100 nM and applied on carbon-coated 400-mesh grids. After blotting, the grids were washed twice with 50 mM Hepes pH 7.5 and stained with 1% uranyl acetate. Data were collected on a 120 kV Philips CM-120 microscope fitted with a Gatan Orius SC1000 detector.

#### Cryo-EM sample preparation

Quantifoil R 2/2 (first dataset) and quantifoil R 0.6/1 (second dataset, Electron Microscopy Sciences) grids were washed with ethyl acetate and glow discharged using a Pelco easiGlow (Pelco) for 60 s at 25 mA (negative polarity). 3 μL of hRFC was applied to a grid at 10 °C and 95% humidity in a Vitrobot Mark IV (FEI). Samples were then blotted at a blotting force of 5 for 5 s after a wait time of 2 s and vitrified by plunging into liquid ethane.

#### Cryo-EM data collection

Grids of hRFC were imaged on a Titan Krios operated at 300 kV. Images were collected on a K3 Summit detector in super-resolution counting mode at a magnification of 81000 ×, with a pixel size of 0.53 Å. The data was collected in two sessions using the multi-hole/multi-shot strategy with SerialEM (Mastronarde 2003) and beam-image shift. During the first session, 3695 micrographs were collected with a target defocus range of −1.2 to −2.3 and using image shift to record three images per hole over four holes with a total dose of 40.3 e−/Å2 per micrograph. During the second session, 7840 micrographs were collected with a target defocus range of −1.2 to −2.6 and using image shift to record one image over four holes with a total dose of 45.0 e−/Å2 per micrograph.

#### Data Processing

The Align Frames module in IMOD (Kremer et al. 1996) was used to align micrograph frames with 2x binning, resulting in a pixel size of 1.06 Å/pixel. For particle picking and 2D classification, dataset two was split into two batches. The micrographs from the different micrograph batches were first processed independently. Initial CTF estimation and particle picking was performed using cisTEM (Rohou and Grigorieff 2015; Grant et al. 2018). Particles were picked with a characteristic radius of 50 Å and a maximum radius of 100 Å. Particles were then extracted with a largest dimension of 170 Å and a box size of 240 pixels and subjected to 2D classification into 50 classes. Particles from classes with well-defined features were extracted for processing in Relion. To increase the number of particles and less well defined side views, the rest of the dataset was subjected to one more round of classification, and more particles that resembled hRFC:PCNA complexes were extracted.

For all further processing steps, all particle stacks were combined yielding 2,933,726 coordinates for putative particles (Supplementary Figure 3). The coordinates and the combined micrographs were imported into Relion 3.0.2 (Zivanov et al. 2018) and CTF parameters were re-estimated with Gctf1.06 (Zhang 2016). Particles were binned to a box size of 120 pixels and 2.12 Å/pixel for the first round of 3D classification. As reference for 3D classification, a refined 3D reconstruction of hRFC bound to PCNA from a preliminary dataset collected on a 200 kV Talos Arctica equipped with a K3 Summit detector was used and down-filtered to 50 Å (Supplemental Figure 3C).

The binned particles were classified into three classes. The 561,558 particles contributing to the best 3D class with well-defined structural features of hRFC and PCNA were extracted without binning. A final round of 3D classification with local angular search helped to obtain a more homogeneous particle stack. These particles were refined to a resolution of 3.5 Å, however some regions (in particular PCNA and the A’ domain) were not well resolved and thus problematic for atomic modeling. We next performed CTF refinement and re-refined the particle stack, which yielded a structure with a reported resolution of 3.4 Å. In this reconstruction however, areas especially within PCNA appeared to suffer from flexibility and misalignment. Suspecting that this was due to motions of PCNA relative to hRFC, we next performed multi-body refinement (Nakane et al. 2018). The best result was obtained with a mask that included the three PCNA-bound and ATPase domains of subunits A-C as well as PCNA (Mask A). Mask B included the collar domain of subunits A-C, as well as complete subunits D and E. After multibody refinement, the density for PCNA and some loops in the A’ domain improved. The reported resolutions for Mask A and Mask B are ∼3.3 Å and ∼3.4 Å, respectively, by FSC gold standard 0.143 criterion (and are ∼3.7 Å and ∼3.8 Å by the FSC 0.5 criterion). The reconstructions of the two Masks were individually sharpened with Relion postprocess and a composite map was generated with the “vop max”function in UCSF Chimera. The composite map was used for atomic model building and refinement. Local resolution estimation using the built-in Relion LocalRes function indicates that loops that locate to the periphery of the complex have lowest resolution and central regions are resolved to better than the FSC-reported resolutions.

#### Model Building

A homology model of the hRFC:PCNA was generated with the structure of yeast RFC (PDB ID: 1SXJ) using Swiss Model (Waterhouse et al. 2018) (Sequence identity between yeast and human subunits: A subunit 24%, B subunit 47%, C subunit 47%, D subunit 53%, E subunit 42%). For PCNA, the human PCNA structure (Gulbis et al. 1996) was used for rigid body fitting. The different subunits were split into individual globular domains that were fit as rigid bodies into the density using UCSF Chimera (Pettersen et al. 2004). The fitted model was carefully adjusted in Coot, and subsequently refined in Phenix with simulated annealing using the real-space refinement tool with rotamer, Ramachandran and secondary structure restraints (Emsley and Cowtan 2004; Liebschner et al. 2019). The refined model was re-examined, re-adjusted in Coot and subjected to further rounds of refinement using the same settings as listed above, but without simulated annealing. UCSF Chimera and Pymol were used for figure generation (Pettersen et al. 2004; DeLano and Others 2002).

## Author Contributions and Notes

C.G., X.L., and B.A.K. designed experiments & analyzed data. C.G. optimized cryo-EM sample preparation, collected and processed data for the preliminary Talos reconstruction. C.G. and X.L. performed further cryo-EM sample preparations, data collection, and data processing. C.G., X.L., J.M., J.L., M.H. expressed and purified constructs and performed biochemical experiments. C.G., X.L.,N.P.S.,and B.A.K. built and refined the atomic model. C.G., X.L., and B.A.K. wrote the paper. The authors declare no conflict of interest.

This article contains supporting information online.

## Acknowledgments

The authors thank Drs. C. Xu, KK Song, and K. Lee for assistance with data collection, and Drs. C. Xu, G. Demo, A. Korostelev, J. Hayes, Mrs. A. Jecrois and Dr. K. Doo Nam for advice on data processing. Furthermore, we would like to thank J. Andrade and Dr. B. Ueberheide for advice with sample preparation for mass spectrometry, sample processing and interpretation of the result. We thank Dr. Stephen J. Benkovic (The Pennsylvania State University) who graciously provided the pCDF-1b vector carrying the RFC1ΔN555 gene and the pET-Duet1 vector carrying the RFC2–3–4–5genes. We thank members of the Kelch, Royer, and Schiffer labs for helpful discussions. This work was funded by the American Cancer Society Research Scholar Award (grant number 440685) and NIGMS (R01-GM127776). C.G. was supported by an Early and Advanced Postdoc Mobility (grant numbers 168972 and 177859) Fellowship of the Swiss National Science Foundation. We thank Emily Agnello and Joshua Pajak for critical reading of the manuscript.

## SUPPLEMENTARY

**Supplemental Table 1: List of BS3 Crosslinks**

**Supplemental Table 2: Structure Determination and Refinement**

**Supplemental Table 3: Somatic cancer mutations found in hRFC**

## Notes

https://www.rcsb.org/structure/6vvo

https://www.ebi.ac.uk/pdbe/entry/emdb/EMD-21405

### Competing Interest Statement

The authors have declared no competing interest.

## References

Abe, Takuya, Masato Ooka, Ryotaro Kawasumi, Keiji Miyata, Minoru Takata, Kouji Hirota, and Dana Branzei. 2018. “Warsaw Breakage Syndrome DDX11 Helicase Acts Jointly with RAD17 in the Repair of Bulky Lesions and Replication through Abasic Sites.” Proceedings of the National Academy of Sciences of the United States of America 115 (33): 8412–17.

Ason, Brandon, Renita Handayani, Christopher R. Williams, Jeffrey G. Bertram, Manju M. Hingorani, Mike O’Donnell, Myron F. Goodman, and Linda B. Bloom. 2003. “Mechanism of Loading the Escherichia Coli DNA Polymerase III *β* Sliding Clamp on DNA: BONA FIDE PRIMER/TEMPLATES PREFERENTIALLY TRIGGER THE *γ* COMPLEX TO HYDROLYZE ATP AND LOAD THE CLAMP.” The Journal of Biological Chemistry 278 (12): 10033–40.

Baple, Emma L., Helen Chambers, Harold E. Cross, Heather Fawcett, Yuka Nakazawa, Barry A. Chioza, Gaurav V. Harlalka, et al. 2014. “Hypomorphic PCNA Mutation Underlies a Human DNA Repair Disorder.” The Journal of Clinical Investigation 124 (7): 3137–46.

Bell, Daphne W., Nilabja Sikdar, Kyoo-Young Lee, Jessica C. Price, Raghunath Chatterjee, Hee-Dong Park, Jennifer Fox, et al. 2011. “Predisposition to Cancer Caused by Genetic and Functional Defects of Mammalian Atad5.” Edited by Nancy Maizels. PLoS Genetics 7 (8): e1002245.

Berdis, Anthony J., and Stephen J. Benkovic. 1996. “Role of Adenosine 5‘-Triphosphate Hydrolysis in the Assembly of the Bacteriophage T4 DNA Replication Holoenzyme Complex.” Biochemistry 35 (28): 9253–65.

Bleichert, Franziska. 2019. “Mechanisms of Replication Origin Licensing: A Structural Perspective.” Current Opinion in Structural Biology 59 (December): 195–204.

Bowman, Gregory D., Mike O’Donnell, and John Kuriyan. 2004. “Structural Analysis of a Eukaryotic Sliding DNA Clamp–clamp Loader Complex.” Nature 429 (6993): 724–30.

Chen, Siying, Mikhail K. Levin, Miho Sakato, Yayan Zhou, and Manju M. Hingorani. 5/2009. “Mechanism of ATP-Driven PCNA Clamp Loading by S. Cerevisiae RFC.” Journal of Molecular Biology 388 (3): 431–42.

Cortese, Andrea, Roberto Simone, Roisin Sullivan, Jana Vandrovcova, Huma Tariq, Wai Yan Yau, Jack Humphrey, et al. 4/2019. “Biallelic Expansion of an Intronic Repeat in RFC1 Is a Common Cause of Late-Onset Ataxia.” Nature Genetics 51 (4): 649–58.

Davey, Megan J., David Jeruzalmi, John Kuriyan, and Mike O’Donnell. 2002. “Motors and Switches: AAA Machines within the Replisome.” Nature Reviews Molecular Cell Biology.

Deimling, F. von, J. M. Scharf, T. Liehr, M. Rothe, A. R. Kelter, P. Albers, W. F. Dietrich, L. M. Kunkel, N. Wernert, and B. Wirth. 1999. “Human and Mouse RAD17 Genes: Identification, Localization, Genomic Structure and Histological Expression Pattern in Normal Testis and Seminoma.” Human Genetics 105 (1-2): 17–27.

DeLano, Warren L., and Others. 2002. “Pymol: An Open-Source Molecular Graphics Tool.” CCP4 Newsletter on Protein Crystallography 40 (1): 82–92.

D’Imprima, Edoardo, Davide Floris, Mirko Joppe, Ricardo Sánchez, Martin Grininger, and Werner Kühlbrandt. 2019. “Protein Denaturation at the Air-Water Interface and How to Prevent It.” eLife 8 (April).

Douzery, Emmanuel J. P., Elizabeth A. Snell, Eric Bapteste, Frédéric Delsuc, and Hervé Philippe. 2004. “The Timing of Eukaryotic Evolution: Does a Relaxed Molecular Clock Reconcile Proteins and Fossils?” Proceedings of the National Academy of Sciences of the United States of America 101 (43): 15386–91.

Duffy, Caroline M., Brendan J. Hilbert, and Brian A. Kelch. 03/2016. “A Disease-Causing Variant in PCNA Disrupts a Promiscuous Protein Binding Site.” Journal of Molecular Biology 428 (6): 1023–40.

Emsley, Paul, and Kevin Cowtan. 2004. “Coot: Model-Building Tools for Molecular Graphics.” Acta Crystallographica Section D Biological Crystallography.

Erzberger, Jan P., and James M. Berger. 2006. “Evolutionary Relationships and Structural Mechanisms of AAA+ Proteins.” Annual Review of Biophysics and Biomolecular Structure 35: 93–114.

Fay, P. J., K. O. Johanson, C. S. McHenry, and R. A. Bambara. 1981. “Size Classes of Products Synthesized Processively by DNA Polymerase III and DNA Polymerase III Holoenzyme of Escherichia Coli.” The Journal of Biological Chemistry 256 (2): 976–83.

Gates, Stephanie N., Adam L. Yokom, Jiabei Lin, Meredith E. Jackrel, Alexandrea N. Rizo, Nathan M. Kendsersky, Courtney E. Buell, et al. 2017. “Ratchet-like Polypeptide Translocation Mechanism of the AAA+ Disaggregase Hsp104.” Science 357 (6348): 273–79.

Gerlach, Piotr, Jan M. Schuller, Fabien Bonneau, Jérôme Basquin, Peter Reichelt, Sebastian Falk, and Elena Conti. 2018. “Distinct and Evolutionary Conserved Structural Features of the Human Nuclear Exosome Complex.” eLife 7 (July).

Goedken, Eric R., Marcia Levitus, Aaron Johnson, Carlos Bustamante, Mike O’Donnell, and John Kuriyan. 3/2004. “Fluorescence Measurements on the E.coli DNA Polymerase Clamp Loader: Implications for Conformational Changes During ATP and Clamp Binding.” Journal of Molecular Biology 336 (5): 1047–59.

Gomes, Xavier V., Sonja L. Gary, and Peter M. J. Burgers. 2000. “Overproduction in Escherichia Coli and Characterization of Yeast Replication Factor C Lacking the Ligase Homology Domain.” The Journal of Biological Chemistry 275 (19): 14541–49.

Gomes, X. V., S. L. G. Schmidt, and P. M. J. Burgers. 2001. “ATP Utilization by Yeast Replication Factor C: II. Multiple Stepwise ATP Binding Events Are Required to Load Proliferating Cell Nuclear Antigen onto Primed DNA.” The Journal of Biological Chemistry 276 (37): 34776–83.

Grant, Timothy, Alexis Rohou, and Nikolaus Grigorieff. 2018. “cisTEM, User-Friendly Software for Single-Particle Image Processing.” eLife. https://doi.org/10.7554/elife.35383.

Gulbis, Jacqueline M., Zvi Kelman, Jerard Hurwitz, Mike O’Donnell, and John Kuriyan. 1996. “Structure of the C-Terminal Region of p21WAF1/CIP1 Complexed with Human PCNA.” Cell 87 (2): 297–306. https://doi.org/10.1016/S0092-8674(00)81347-1.

Hedglin, Mark, Senthil K. Perumal, Zhenxin Hu, and Stephen Benkovic. 2013. “Stepwise Assembly of the Human Replicative Polymerase Holoenzyme.”. eLife.

Hingorani, Manju M., and Mike O’Donnell. 1998. “ATP Binding to theEscherichia coliClamp Loader Powers Opening of the Ring-Shaped Clamp of DNA Polymerase III Holoenzyme.” Journal of Biological Chemistry.

Huang, C. C., J. E. Hearst, and B. M. Alberts. 1981. “Two Types of Replication Proteins Increase the Rate at Which T4 DNA Polymerase Traverses the Helical Regions in a Single-Stranded DNA Template.” The Journal of Biological Chemistry 256 (8): 4087–94.

Jarvis, T. C., L. S. Paul, J. W. Hockensmith, and P. H. von Hippel. 1989. “Structural and Enzymatic Studies of the T4 DNA Replication System. II. ATPase Properties of the Polymerase Accessory Protein Complex.” The Journal of Biological Chemistry 264 (21): 12717–29.

Johnson, Aaron, Nina Y. Yao, Gregory D. Bowman, John Kuriyan, and Mike O’Donnell. 2006. “The Replication Factor C Clamp Loader Requires Arginine Finger Sensors to Drive DNA Binding and Proliferating Cell Nuclear Antigen Loading.” The Journal of Biological Chemistry 281 (46): 35531–43.

Kadyrov, Farid A., Jochen Genschel, Yanan Fang, Elisabeth Penland, Winfried Edelmann, and Paul Modrich. 2009. “A Possible Mechanism for Exonuclease 1-Independent Eukaryotic Mismatch Repair.” Proceedings of the National Academy of Sciences of the United States of America 106 (21): 8495–8500.

Kazmirski, Steven L., Marjetka Podobnik, Tanya F. Weitze, Mike O’Donnell, and John Kuriyan. 2004. “Structural Analysis of the Inactive State of the Escherichia Coli DNA Polymerase Clamp-Loader Complex.” Proceedings of the National Academy of Sciences of the United States of America 101 (48): 16750–55.

Kelch, Brian A. 08/2016. “Review: The Lord of the Rings: Structure and Mechanism of the Sliding Clamp Loader: The Lord of the Rings.” Edited by Alfred Wittinghofer. Biopolymers 105 (8): 532–46.

Kelch, Brian A., Debora L. Makino, Mike O’Donnell, and John Kuriyan. 2011. “How a DNA Polymerase Clamp Loader Opens a Sliding Clamp.” Science 334 (6063): 1675–80.

Kelch, Brian A., Debora L. Makino, Mike O’Donnell, and John Kuriyan. 2012. “Clamp Loader ATPases and the Evolution of DNA Replication Machinery.” BMC Biology 10 (1): 34.

Kremer, J. R., D. N. Mastronarde, and J. R. McIntosh. 1996. “Computer Visualization of Three-Dimensional Image Data Using IMOD.” Journal of Structural Biology 116 (1): 71–76.

Kroker, Alice J., and John B. Bruning. 2015. “p21 Exploits Residue Tyr151 as a Tether for High-Affinity PCNA Binding.” Biochemistry 54 (22): 3483–93.

Langston, Lance D., and Mike O’Donnell. 2008. “DNA Polymerase *δ* Is Highly Processive with Proliferating Cell Nuclear Antigen and Undergoes Collision Release upon Completing DNA.” The Journal of Biological Chemistry 283 (43): 29522–31.

Leu, Frank P., Manju M. Hingorani, Jennifer Turner, and Mike O’Donnell. 2000. “The *δ* Subunit of DNA Polymerase III Holoenzyme Serves as a Sliding Clamp Unloader in Escherichia Coli.” The Journal of Biological Chemistry 275 (44): 34609–18.

Liebschner, Dorothee, Pavel V. Afonine, Matthew L. Baker, Gábor Bunkóczi, Vincent B. Chen, Tristan I. Croll, Bradley Hintze, et al. 2019. “Macromolecular Structure Determination Using X-Rays, Neutrons and Electrons: Recent Developments in Phenix.” Acta Crystallographica. Section D, Structural Biology 75 (Pt 10): 861–77.

Liu, Huanting, and James H. Naismith. 2008. “An Efficient One-Step Site-Directed Deletion, Insertion, Single and Multiple-Site Plasmid Mutagenesis Protocol.” BMC Biotechnology 8 (1): 91.

Liu, Juan, Yayan Zhou, and Manju M. Hingorani. 2017. “Linchpin DNA-Binding Residues Serve as Go/no-Go Controls in the Replication Factor C-Catalyzed Clamp Loading Mechanism.” The Journal of Biological Chemistry, August, jbc.M117.798702.

Li, Yanling, Sijie Gan, Lin Ren, Long Yuan, Junlan Liu, Wei Wang, Xiaoyu Wang, et al. 2018. “Multifaceted Regulation and Functions of Replication Factor C Family in Human Cancers.” American Journal of Cancer Research 8 (8): 1343–55.

Majka, Jerzy, and Peter M. J. Burgers. 2004. “The PCNA–RFC Families of DNA Clamps and Clamp Loaders.” Progress in Nucleic Acid Research and Molecular Biology 78: 227–60.

Maki, S., and A. Kornberg. 1988. “DNA Polymerase III Holoenzyme of Escherichia Coli. II. A Novel Complex Including the Gamma Subunit Essential for Processive Synthesis.” The Journal of Biological Chemistry 263 (14): 6555–60.

Mastronarde, David N. 2003. “SerialEM: A Program for Automated Tilt Series Acquisition on Tecnai Microscopes Using Prediction of Specimen Position.” Microscopy and Microanalysis: The Official Journal of Microscopy Society of America, Microbeam Analysis Society, Microscopical Society of Canada 9 (S02): 1182–83.

Moldovan, George-Lucian, Boris Pfander, and Stefan Jentsch. 05/2007. “PCNA, the Maestro of the Replication Fork.” Cell 129 (4): 665–79.

Mondol, Tanumoy, Joseph L. Stodola, Roberto Galletto, and Peter M. Burgers. 2019. “PCNA Accelerates the Nucleotide Incorporation Rate by DNA Polymerase *δ*.” Nucleic Acids Research 47 (4): 1977–86.

Nakane, Takanori, Dari Kimanius, Erik Lindahl, and Sjors Hw Scheres. 2018. “Characterisation of Molecular Motions in Cryo-EM Single-Particle Data by Multi-Body Refinement in RELION.” eLife 7 (June).

Noble, Alex J., Hui Wei, Venkata P. Dandey, Zhening Zhang, Yong Zi Tan, Clinton S. Potter, and Bridget Carragher. 10/2018. “Reducing Effects of Particle Adsorption to the Air–water Interface in Cryo-EM.” Nature Methods 15 (10): 793–95.

Ochoa, David, Andrew F. Jarnuczak, Cristina Viéitez, Maja Gehre, Margaret Soucheray, André Mateus, Askar A. Kleefeldt, et al. 2019. “The Functional Landscape of the Human Phosphoproteome.” Nature Biotechnology, December.

Peled, Michael, Anna S. Tocheva, Sabina Sandigursky, Shruti Nayak, Elliot A. Philips, Kim E. Nichols, Marianne Strazza, et al. 2018. “Affinity Purification Mass Spectrometry Analysis of PD-1 Uncovers SAP as a New Checkpoint Inhibitor.” Proceedings of the National Academy of Sciences of the United States of America 115 (3): E468–77.

Perumal, Senthil K., Xiaojun Xu, Chunli Yan, Ivaylo Ivanov, and Stephen J. Benkovic. 2019. “Recognition of a Key Anchor Residue by a Conserved Hydrophobic Pocket Ensures Subunit Interface Integrity in DNA Clamps.” Journal of Molecular Biology 431 (14): 2493–2510.

Pettersen, Eric F., Thomas D. Goddard, Conrad C. Huang, Gregory S. Couch, Daniel M. Greenblatt, Elaine C. Meng, and Thomas E. Ferrin. 2004. “UCSF Chimera—a Visualization System for Exploratory Research and Analysis.” Journal of Computational Chemistry 25 (13): 1605–12.

Pietroni, Paola, Mark C. Young, Gary J. Latham, and Peter H. von Hippel. 1997. “Structural Analyses of gp45 Sliding Clamp Interactions during Assembly of the Bacteriophage T4 DNA Polymerase Holoenzyme I. Conformational Changes within the gp44/62-gp45-ATP Complex during Clamp Loading.” The Journal of Biological Chemistry 272 (50): 31666–76.

Podust, Vladimir N., Nikhil Tiwari, Scott Stephan, and Ellen Fanning. 1998. “Replication Factor C Disengages from Proliferating Cell Nuclear Antigen (PCNA) upon Sliding Clamp Formation, and PCNA Itself Tethers DNA Polymerase *δ* to DNA.” The Journal of Biological Chemistry 273 (48): 31992–99.

Ponzoni, Luca, Daniel A. Peñaherrera, Zoltán N. Oltvai, and Ivet Bahar. 2020. “Rhapsody: Predicting the Pathogenicity of Human Missense Variants.” Bioinformatics, February.

Puchades, Cristina, Anthony J. Rampello, Mia Shin, Christopher J. Giuliano, R. Luke Wiseman, Steven E. Glynn, and Gabriel C. Lander. 2017. “Structure of the Mitochondrial Inner Membrane AAA+ Protease YME1 Gives Insight into Substrate Processing.” Science 358 (6363).

Puchades, Cristina, Colby R. Sandate, and Gabriel C. Lander. 2020. “The Molecular Principles Governing the Activity and Functional Diversity of AAA+ Proteins.” Nature Reviews. Molecular Cell Biology 21 (1): 43–58.

Rohou, Alexis, and Nikolaus Grigorieff. 2015. “CTFFIND4: Fast and Accurate Defocus Estimation from Electron Micrographs.” Journal of Structural Biology 192 (2): 216–21.

Rozbeský, Daniel, Michal Rosůlek, Zdeněk Kukačka, Josef Chmelík, Petr Man, and Petr Novák. 2018. “Impact of Chemical Cross-Linking on Protein Structure and Function.” Analytical Chemistry 90 (2): 1104–13.

Sakato, Miho, Mike O’Donnell, and Manju M. Hingorani. 02/2012. “A Central Swivel Point in the RFC Clamp Loader Controls PCNA Opening and Loading on DNA.” Journal of Molecular Biology 416 (2): 163–75.

Schmidt, Sonja L. Gary, Sonja L. Gary Schmidt, Angela L. Pautz, and Peter M. J. Burgers. 2001. “ATP Utilization by Yeast Replication Factor C.” Journal of Biological Chemistry.

Seybert, Anja, and Dale B. Wigley. 2004. “Distinct Roles for ATP Binding and Hydrolysis at Individual Subunits of an Archaeal Clamp Loader.” The EMBO Journal 23 (6): 1360–71.

Simonetta, Kyle R., Steven L. Kazmirski, Eric R. Goedken, Aaron J. Cantor, Brian A. Kelch, Randall McNally, Steven N. Seyedin, Debora Makino, Mike O’Donnell, and John Kuriyan. 05/2009. “The Mechanism of ATP-Dependent Primer-Template Recognition by a Clamp Loader Complex.” Cell 137 (4): 659–71.

Stodola, Joseph L., and Peter M. Burgers. 2016-5. “Resolving Individual Steps of Okazaki Fragment Maturation at Msec Time-Scale.” Nature Structural & Molecular Biology 23 (5): 402–8.

Sun, Qiming, Toshiki Tsurimoto, Franceline Juillard, Lin Li, Shijun Li, Erika De León Vázquez, She Chen, and Kenneth Kaye. 2014. “Kaposi’s Sarcoma-Associated Herpesvirus LANA Recruits the DNA Polymerase Clamp Loader to Mediate Efficient Replication and Virus Persistence.” Proceedings of the National Academy of Sciences of the United States of America 111 (32): 11816–21.

Tainer, John A., J. Andrew McCammon, and Ivaylo Ivanov. 2010. “Recognition of the Ring-Opened State of Proliferating Cell Nuclear Antigen by Replication Factor C Promotes Eukaryotic Clamp-Loading.” Journal of the American Chemical Society 132 (21): 7372–78.

Tang, Hui, Benjamin Hilton, Phillip R. Musich, Ding Zhi Fang, and Yue Zou. 04/2012. “Replication Factor C1, the Large Subunit of Replication Factor C, Is Proteolytically Truncated in Hutchinson-Gilford Progeria Syndrome: Truncation of Replication Factor C1 in HGPS.” Aging Cell 11 (2): 363–65.

Trakselis, M. A., S. C. Alley, E. Abel-Santos, and S. J. Benkovic. 2001. “Creating a Dynamic Picture of the Sliding Clamp during T4 DNA Polymerase Holoenzyme Assembly by Using Fluorescence Resonance Energy Transfer.” Proceedings of the National Academy of Sciences 98 (15): 8368–75.

Turner, Jennifer, Manju M. Hingorani, Zvi Kelman, and Mike O’Donnell. 1999. “The Internal Workings of a DNA Polymerase Clamp-Loading Machine.” The EMBO Journal 18 (3): 771–83.

Uhlmann, Frank, Jinsong Cai, Emma Gibbs, Mike O’Donnell, and Jerard Hurwitz. 1997. “Deletion Analysis of the Large Subunit p140 in Human Replication Factor C Reveals Regions Required for Complex Formation and Replication Activities.” The Journal of Biological Chemistry 272 (15): 10058–64.

Wang, Shao-Chun. 2014. “PCNA: A Silent Housekeeper or a Potential Therapeutic Target?” Trends in Pharmacological Sciences 35 (4): 178–86.

Wang, Xin, Christine M. Helfer, Neha Pancholi, James E. Bradner, and Jianxin You. 2013. “Recruitment of Brd4 to the Human Papillomavirus Type 16 DNA Replication Complex Is Essential for Replication of Viral DNA.” Journal of Virology 87 (7): 3871–84.

Wang, Xin, Jing Li, Rachel M. Schowalter, Jing Jiao, Christopher B. Buck, and Jianxin You. 2012. “Bromodomain Protein Brd4 Plays a Key Role in Merkel Cell Polyomavirus DNA Replication.” PLoS Pathogens 8 (11): e1003021.

Waterhouse, Andrew, Martino Bertoni, Stefan Bienert, Gabriel Studer, Gerardo Tauriello, Rafal Gumienny, Florian T. Heer, et al. 2018. “SWISS-MODEL: Homology Modelling of Protein Structures and Complexes.” Nucleic Acids Research 46 (W1): W296–303.

Wilson, Rosemary H. C., Antonio J. Biasutto, Lihao Wang, Roman Fischer, Emma L. Baple, Andrew H. Crosby, Erika J. Mancini, and Catherine M. Green. 02/2017. “PCNA Dependent Cellular Activities Tolerate Dramatic Perturbations in PCNA Client Interactions.” DNA Repair 50: 22–35.

Yoo, Jiho, Mengyu Wu, Ying Yin, Mark A. Herzik, Gabriel C. Lander, and Seok-Yong Lee. 2018. “Cryo-EM Structure of a Mitochondrial Calcium Uniporter.” Science, June, eaar4056.

Zhang, Kai. 2016. “Gctf: Real-Time CTF Determination and Correction.” Journal of Structural Biology 193 (1): 1–12.

Zhou, Yayan, and Manju M. Hingorani. 2012. “Impact of Individual Proliferating Cell Nuclear Antigen-DNA Contacts on Clamp Loading and Function on DNA.” The Journal of Biological Chemistry 287 (42): 35370–81.

Zivanov, Jasenko, Takanori Nakane, Björn O. Forsberg, Dari Kimanius, Wim Jh Hagen, Erik Lindahl, and Sjors Hw Scheres. 2018. “New Tools for Automated High-Resolution Cryo-EM Structure Determination in RELION-3.” eLife 7 (November).

