## Supplemental Information and Supplemental Table 2 for "Structure of the human clamp loader bound to the sliding clamp: a further twist on AAA+ mechanism"

**Supplemental Table 2: Structure determination and refinement.**

| **Deposited structures** | |
|---|---|
| PDB accession no. | 6VVO |
| EMDB accession no. | 21405 |

| **Data collection** | |
|---|---|
| Microscope | FEI Titan Krios |
| Detector | Gatan K-3 |
| Voltage (kV) | 300 |
| Magnification | 81,000 |
| Electron exposure ($e^-$/Å$^2$) | 40 (dataset 1) /45 (dataset 2) |
| Defocus range (µm) | -1,2 to -2,6 |
| Pixel size (Å) | 0.53 |
| Number of Micrographs | 11,535 |

| **Data processing** | |
|---|---|
| Initial number of particles | 2,933,726 |
| Final number of particles | 193,943 |
| Map-sharpening B-factor (Å$^2$) | -140 (Mask A) / -152 (Mask B) |
| Final resolution (Å) | 3.3 (Mask A) / 3.4 (Mask B) |

| **Asymmetric unit refinement** | |
|---|---|
| Map correlation (%) | 85 |
| R.M.S.D. (bonds) | 0.008 |
| R.M.S.D. (angles) | 0.816 |
| MolProbity score | 1.85 |
| All-atom clashscore | 9.76 |
| Ramachandran favored (%) | 95.16 |
| Ramachandran allowed (%) | 4.72 |
| Ramachandran Outliers (%) | 0.12 |
| Rotamer outliers (%) | 0.41 |
| C-beta deviations | 0 |

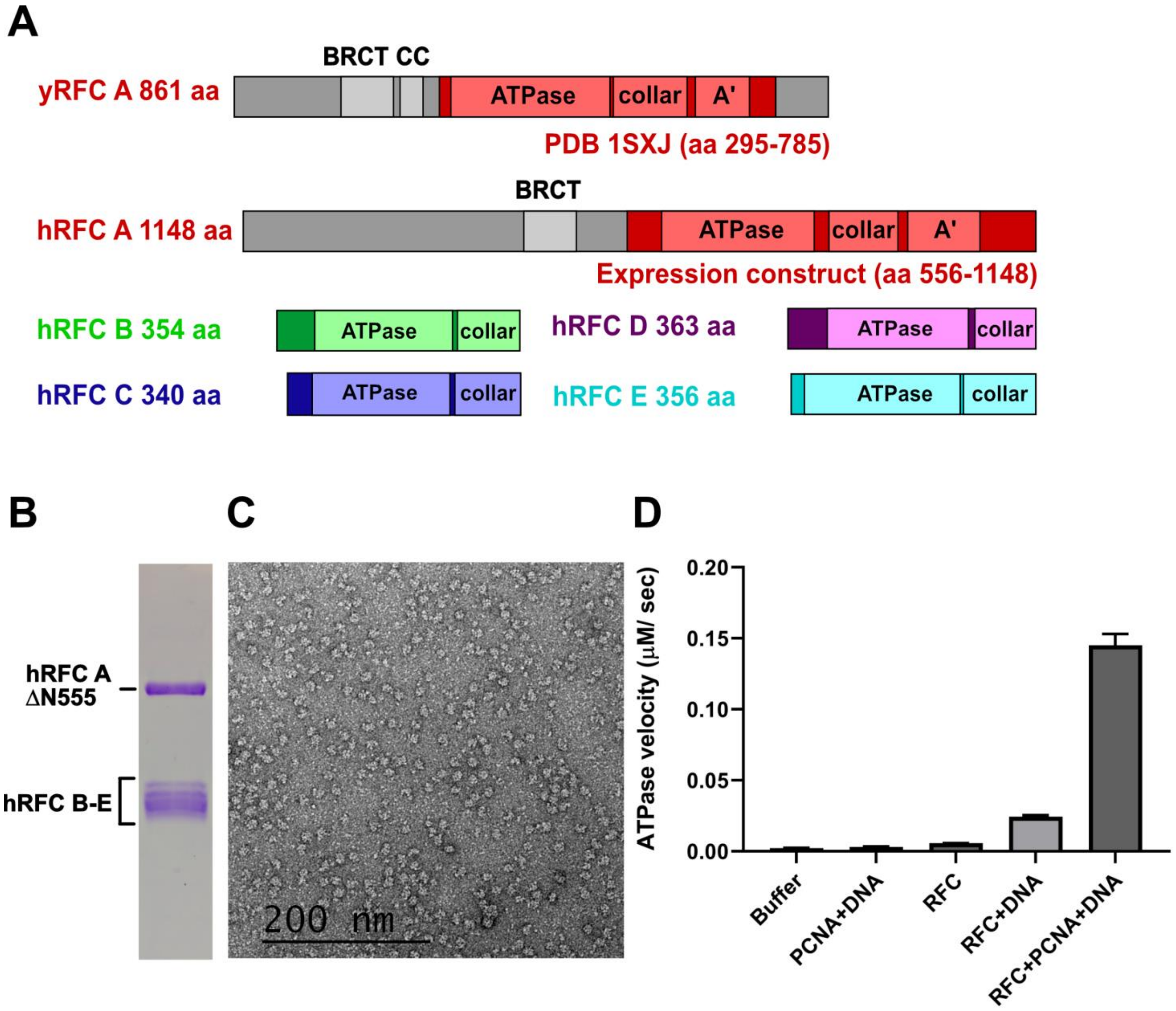


**Supplementary Figure 1. Purification and Characterization of hRFC (Rfc1ΔN555)**

**(A)** Color-coded primary sequence of human RFC complex shows the domain organization and expression construct design. Greyed regions of the protein are not included in the expression construct. For comparison, the sequence of *S.cerevisiae* Rfc1 that was visible in the crystal structure of yeast RFC is shown (PDB 1SXJ). **(B)** SDS-PAGE gel of purified hRFC (RFC1ΔN555). **(C)** Negative stain of hRFC (RFC1ΔN555) shows homogenous and evenly dispersed particles. **(D)** ATPase activity of hRFC (RFC1ΔN555) .

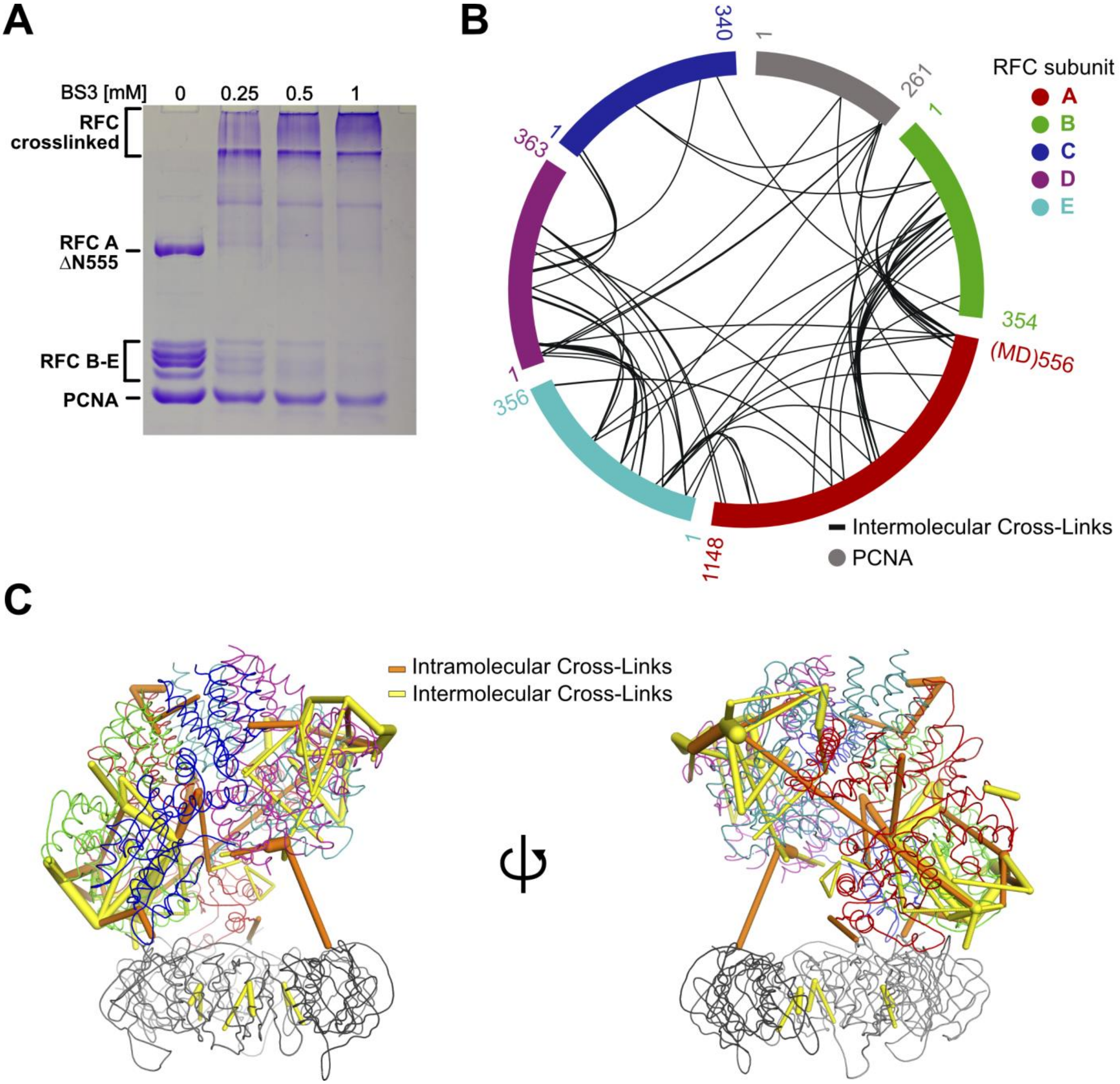


**Supplementary Figure 2. Crosslinking-Mass Spectrometry of the hRFC (RFC1ΔN555) : PCNA complex.** **(A)** Titration of BS3 (bis(sulfosuccinimidyl)suberate). At a concentration of 1 mM the majority of hRFC, but not all PCNA, is cross-linked. **(B)** Schematic representation of the 76 interprotein crosslinks identified in hRFC (RFC1ΔN555). A represents RFC1ΔN555, B RFC2, C RFC5, D RFC4, and E RFC3. The C-terminus of the A subunit starts at amino acid 556 of RFC1 and is preceded by Methionine and an Aspartate (MD) in this construct. Crosslinked residues are listed in Supplementary Table 1. **(C)** Close-up on the cross-links that were mapped on the structure of hRFC(RFC1ΔN555). All intramolecular crosslinks that could be mapped on the structure are shown (51 of 97 total). For the display of the intermolecular crosslinks, a cutoff of a minimal cross-link score of 57.6 was applied.

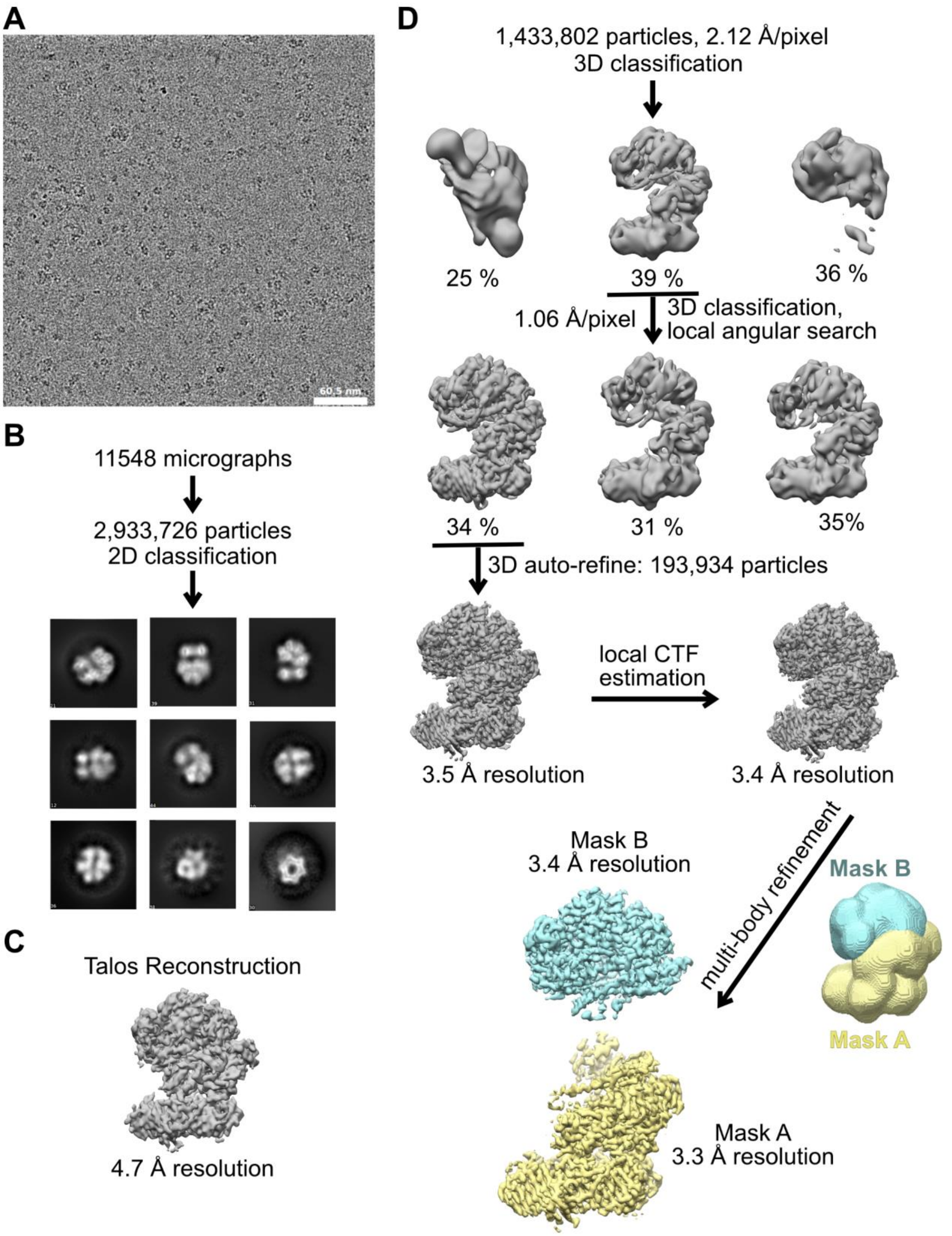


**Supplementary Figure 3. Schematic of hRFC:PCNA cryo-EM refinement and classification.**
**(A)** Micrographs were taken on a Thermo Fisher Scientific Titan Krios with a Gatan K3 detector. **(B)** 2D classification shows different views of the complex. **(C)** A dataset collected on a Talos Arctica with Gatan K3 detector was processed with cisTEM and Relion to generate an initial *ab intio* reconstruction of the complex. The Talos reconstruction and the Krios reconstruction are in the same conformation. **(D)** The first round of 3D classification was performed with the 2x binned particle stack and the Talos reconstruction filtered to 50 Å as reference. A further round of classification, refinement after local CTF estimation and multibody refinement further improved the resolution.

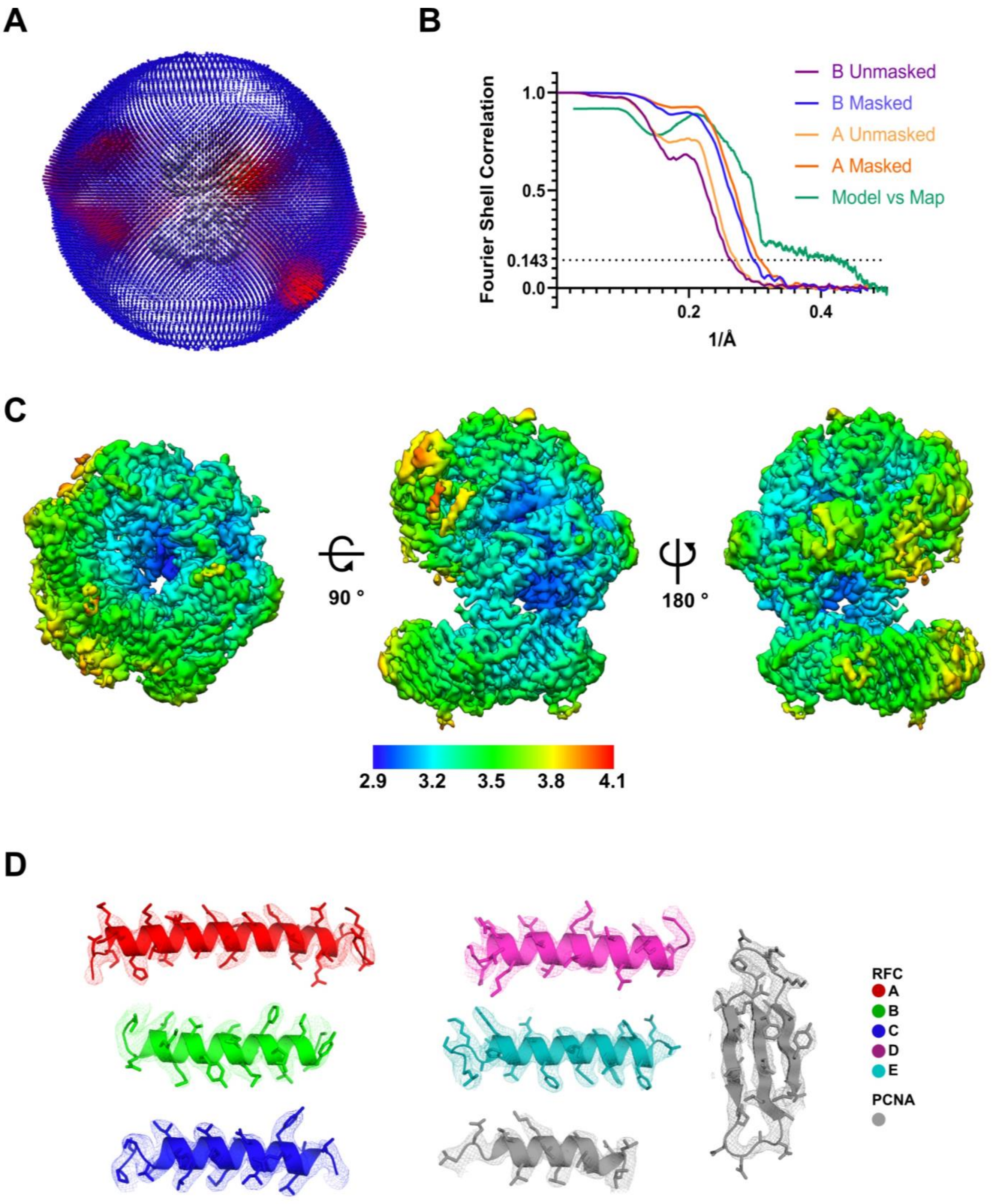


**Supplementary Figure 4. Quality of hRFC:PCNA map and model.**
**(A)** Angular distribution of particles used for the final reconstruction. **(B)** Fourier shell correlation (FSC) curves for the two halves of the hRFC:PCNA multibody reconstruction; resolution is 3.3 Å (Mask A) and 3.4 Å (Mask B) at the gold standard FSC cutoff of 0.143, together with model vs map FSC curve for the whole complex. **(C)** Local resolution of the hRFC:PCNA reconstruction. **(D)** Representative sections of each complex subunit of the density map with fitted model.

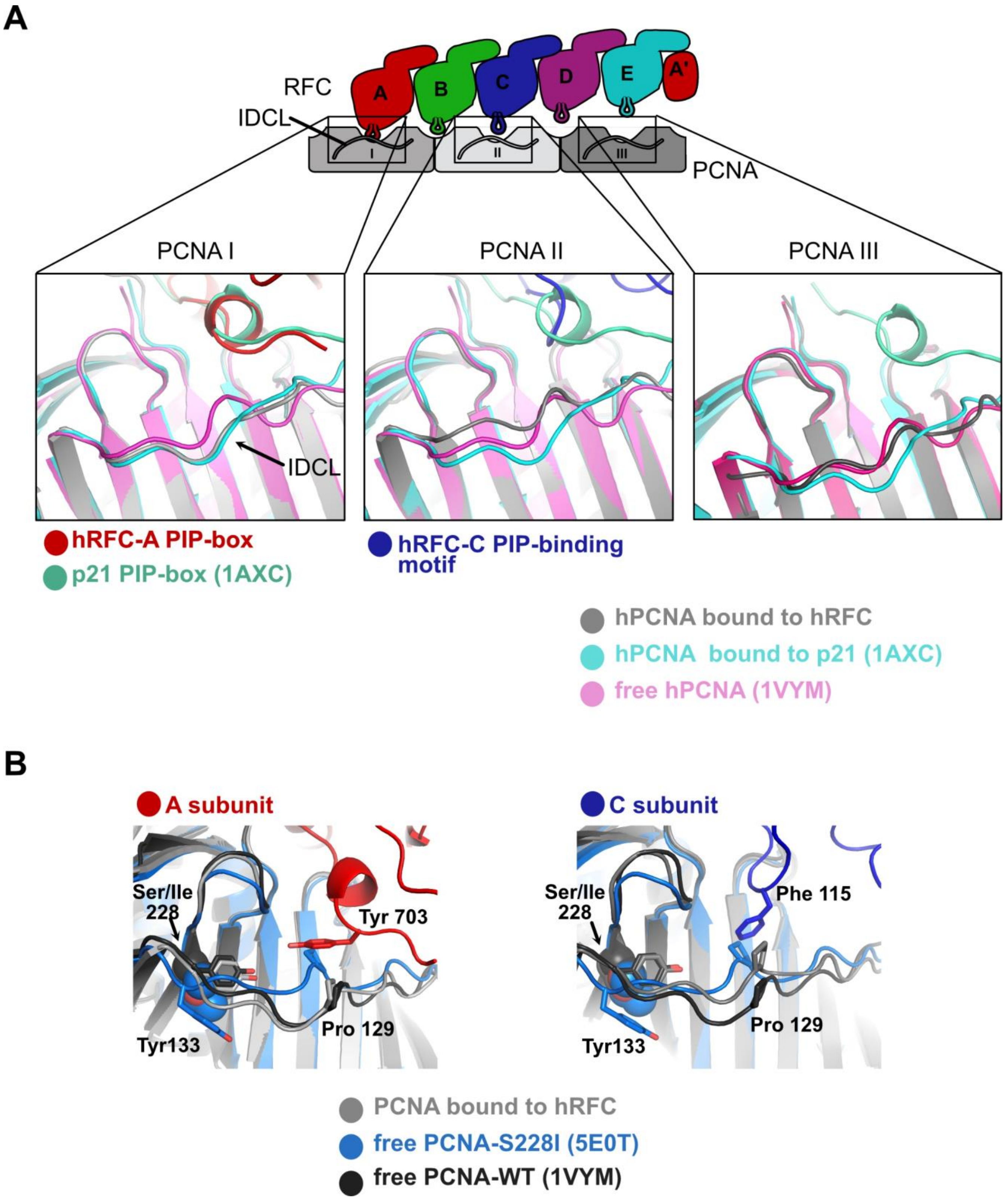


**Supplementary Figure 5. PCNA inter domain connecting loop (IDCL) conformation.**
**(A)** The IDCL adopts different conformations upon interaction with the A and C subunits of hRFC. **(B)** Comparison of the IDCL of PCNA subunits I and II with the IDCL of free S228I-PCNA and wild-type PCNA. Free S288I-PCNA is sterically unable to bind the A- or C-subunit without a conformational change (PCNA-Pro129 would clash with A-Tyr 703 and C-Phe115, respectively).

**A**

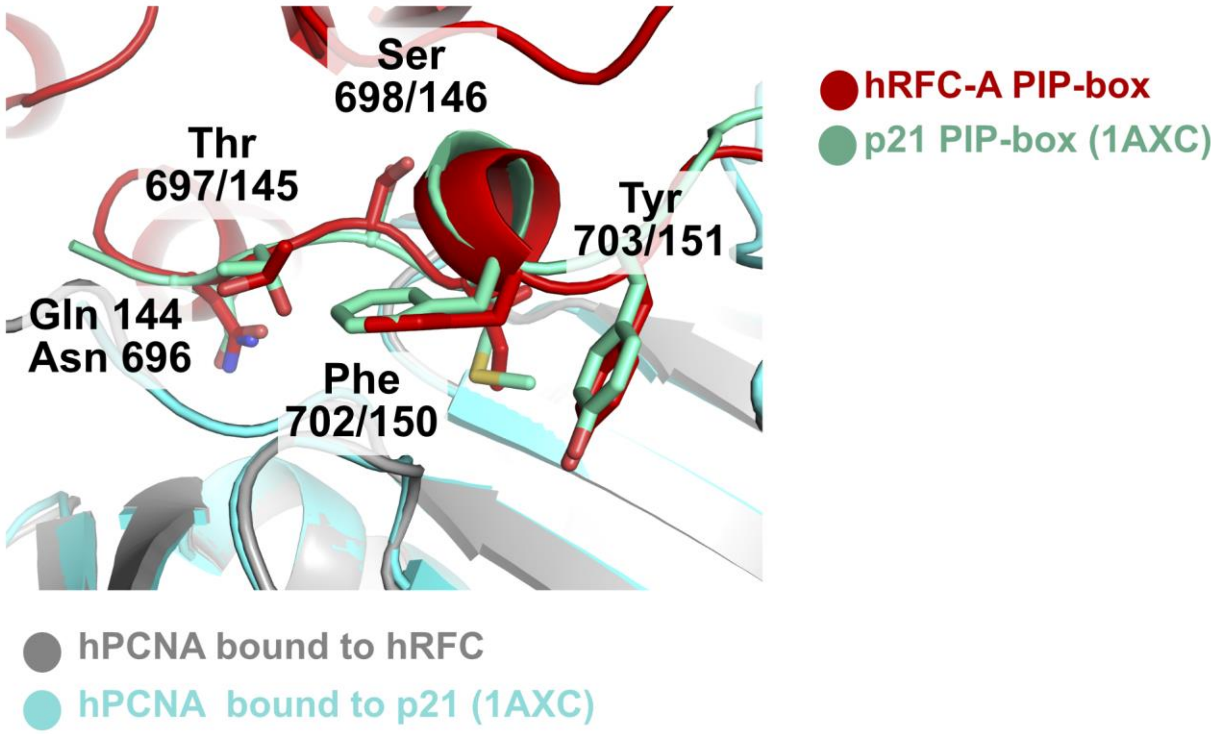


**B**

```
p21        143   RQTSMTDFY 151
                 ••||•   ||
RFC1(A)    695   NNTSIKGFY 703
Consensus        xQxxΨxxΩΩ
                   Ψ=M,I,L,V
                   Ω=F,Y
```

**Supplementary Figure 6. Similar PCNA binding modes by the hRFC A subunit and p21.**
**(A)** Comparison of the hRFC A-subunit PIP-box with the p21 PIP-box. **(B)** Sequence alignment of the primary PCNA binding regions for p21 and RFC1.

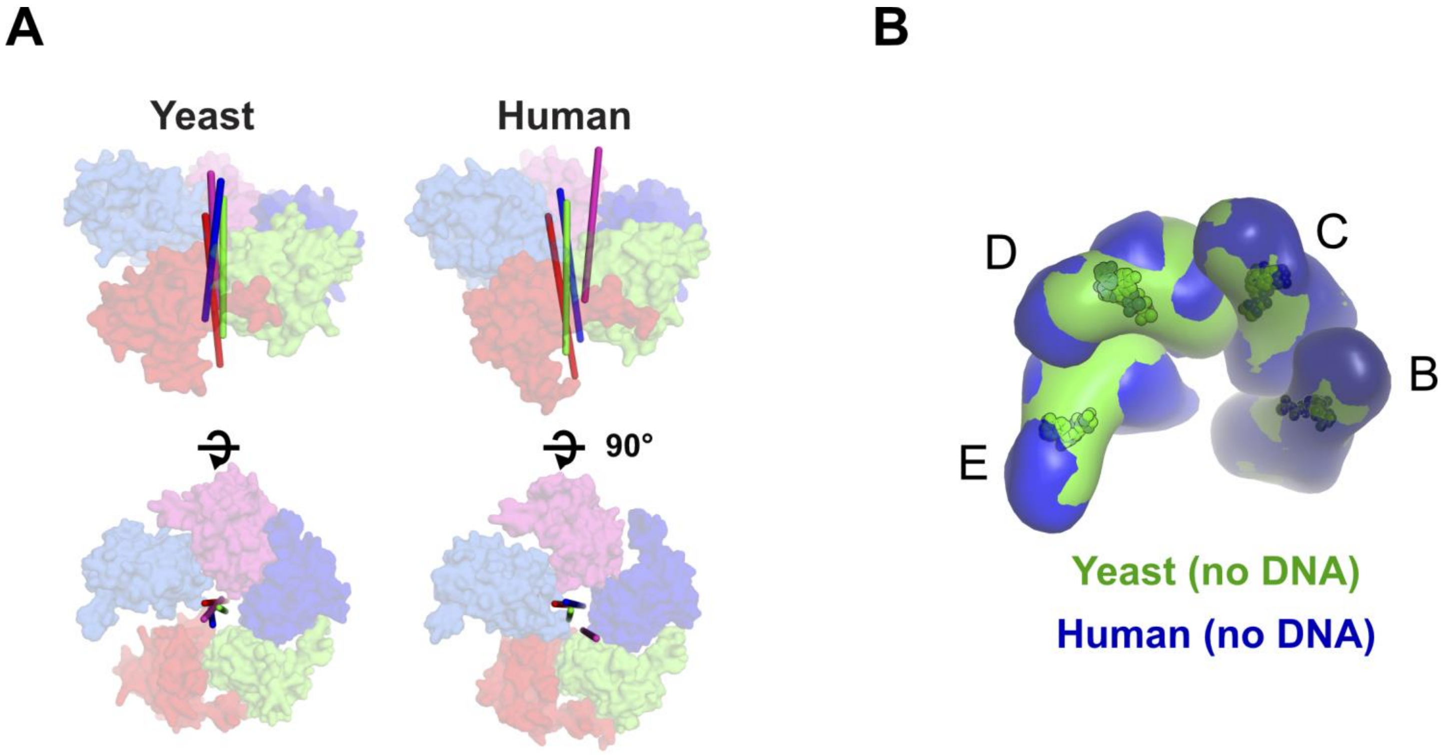


**Supplementary Figure 7. Architecture of the AAA-ATP domains and nucleotide binding.**
**(A)** Diagrams show the side view (top panel), and the top view (middle) of the Rossman fold domains of yeast RFC (PDB 1SXJ), and hRFC. The rotation axes that relate the A to B, the B to C, the C to D and the D to E subunits are shown in red, green, blue, and magenta, respectively. The rotation axes of yeast are mildly skewed; the rotation axes of hRFC are severely skewed. **(B)** The AAA+ spirals of both yRFC and hRFC are overtwisted.

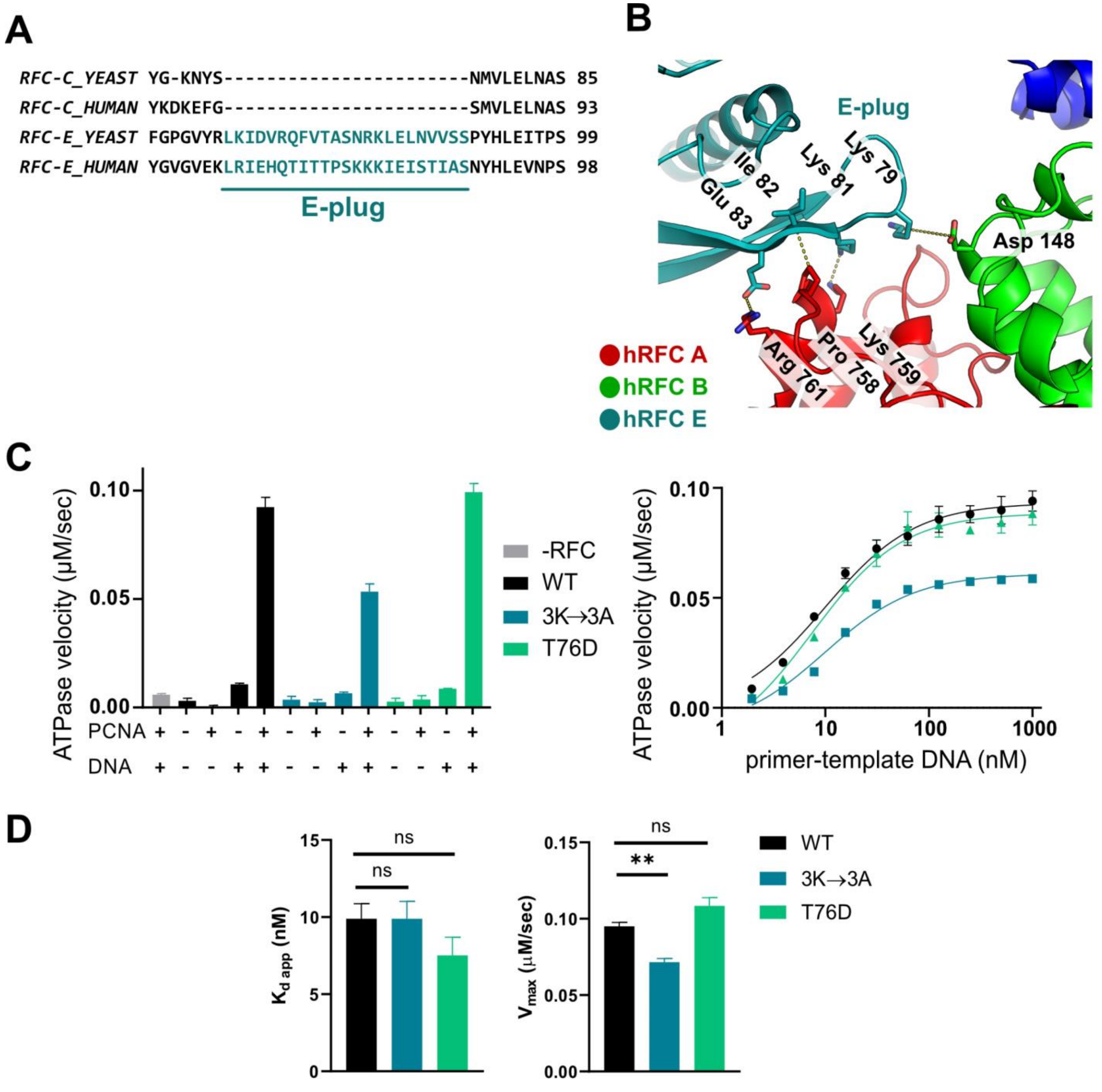


| | WT | 3K→3A | T76D |
|---|---|---|---|
| $K_{d\ app}$ (nM) | 9.9±1.0 | 9.9±1.1 | 7.5±1.2 |
| $V_{max}$ (μM/s) | 0.095±0.002 | 0.072±0.002 | 0.108±0.005 |

**Supplementary Figure 8. E-plug conservation, interactions, and effect of 'E-plug' mutants on DNA binding and activity.**

**(A)** The E-plug is conserved in RFC E subunits but is not found in other subunits. Sequence alignment shows the conservation of the E-plug between *S. cerevisiae* (YEAST) and *H. sapiens* (HUMAN). **(B)** The E-plug blocks the A-gate by binding directly to the Rossmann fold of the A subunit, and possibly the B subunit. Several residues mediating this interaction are highlighted. Although the sidechain position of Lys79-E is ambiguous, it is possible to make an electrostatic interaction with Asp148-B. **(C)** ATPase activity and DNA binding affinity of hRFC 'E-plug' mutants. **(D)** Both 'E-plug' mutants bind DNA with similar affinity than WT-hRFC (left). The maximum ATP hydrolysis rate of the 3K → 3A mutant is significantly reduced (P value = 0.0083, right panel).

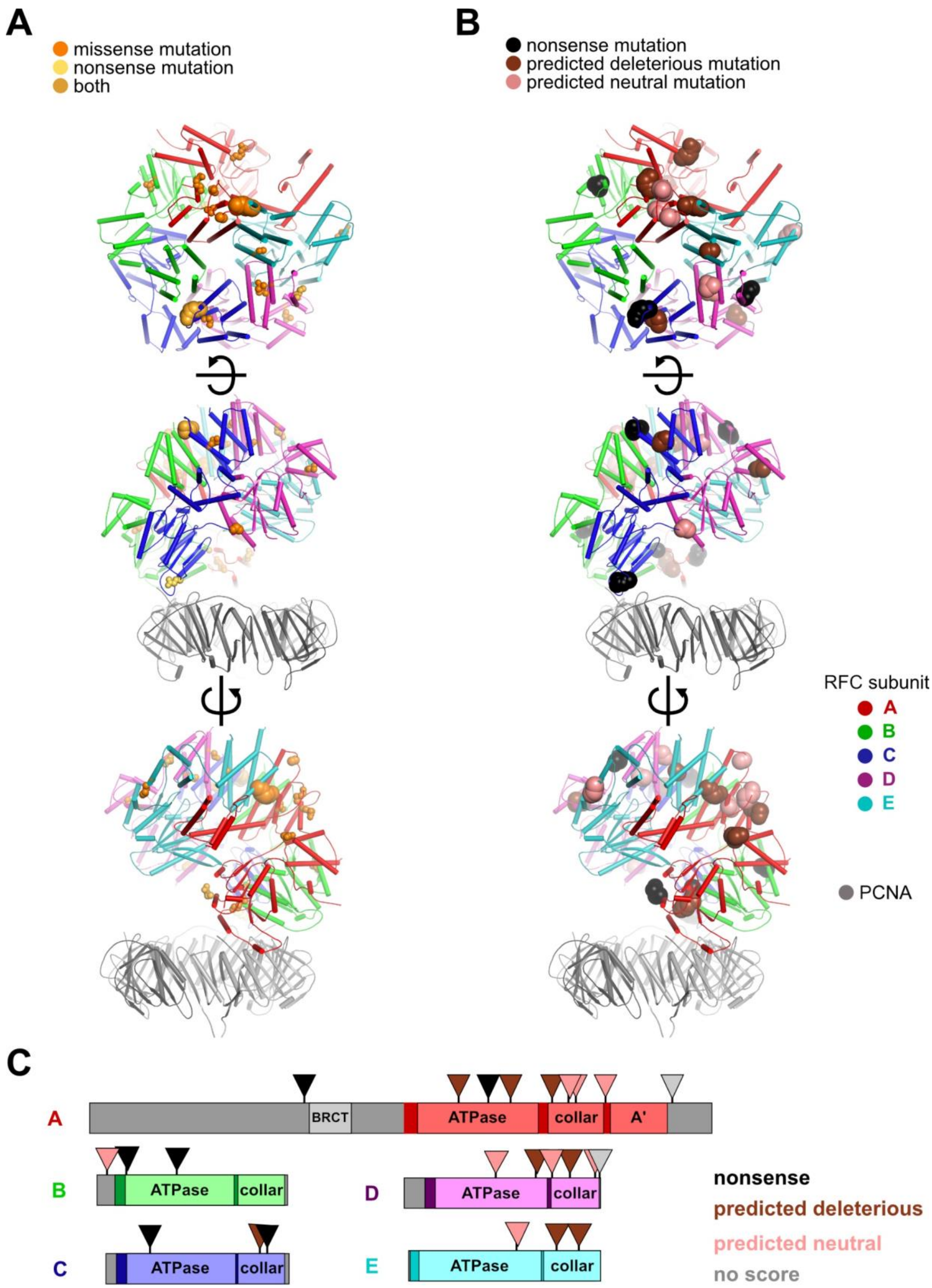


**Supplementary Figure 9. Mapping of cancer somatic mutations on hRFC.**
Cancer mutations from the COSMIC database are mapped onto the hRFC:PCNA structure. **(A)** The size of the spheres are scaled by the number of hits in the COSMIC at that particular position, so that sites with numerous hits are displayed larger. Sites are also color-coded according to the type of mutations observed. **(B)** Missense mutation variants are color-coded according to with Rhapsody predicted pathogenicity and mapped onto the 3D structure and **(C)** the primary sequence of hRFC:PCNA .
